# The development of coarse and fine feature tuning in infant visual cortex

**DOI:** 10.64898/2026.10.14.758935

**Authors:** Áine T. Dineen, Clíona O’Doherty, Anna Truzzi, Anna Kravchenko, Alex R. Wade, Graham King, Enna-Louise D’Arcy, Angela T. Byrne, Ailbhe Tarrant, Adrienne Foran, Eleanor J. Molloy, Rhodri Cusack

## Abstract

Objects can be recognized from visual features at multiple spatial scales, from coarse global shape to fine local detail. How infant visual cortex encodes these features across the visual hierarchy, however, remains unknown. We combined awake infant fMRI with developmentally inspired deep neural networks (DNNs) trained and tested with different levels of image blur to probe cortical tuning to coarse and fine visual features. V1 showed mature tuning to coarse and fine features by 2 months. In contrast, infant extrastriate cortex (V2–V4) followed a protracted developmental trajectory and was biased toward coarse object features relative to adults. This relative bias persisted at 9 months and was not readily explained by developing acuity. The sustained coarse-feature bias contrasted with DNNs, in which low-spatial-frequency representations took longest to converge with visual cortex during training, revealing distinct developmental biases in object-feature tuning in biological and artificial visual systems.

**Highlights:** • Blur-trained DNNs reveal coarse and fine feature tuning in visual cortex.

• V1 shows mature tuning to coarse and fine visual features by 2 months.

• Extrastriate tuning remains immature and relatively coarse-biased through 9 months.

• This coarse bias is not readily explained by developing acuity.

• In DNNs, coarse features take longest to converge with visual cortex during training.

## Introduction

When an infant sees an object, what features does their visual cortex encode? Objects contain information at multiple spatial scales, from fine-grained texture to more spatially extended shape features. A cat, for example, has the coarse shape of its head and body alongside the fine texture of its fur. In adults, these features are represented along the ventral visual stream (VVS), which progresses from relatively simple perceptual features in early visual areas, through contours and textures, to increasingly abstract object representations such as category.^1–4^ How tuning to coarse- and fine-scale object features develops along the infant VVS remains poorly understood.^5^

Early infant vision is strikingly coarse. Visual acuity, the ability to resolve fine spatial detail, and contrast sensitivity, the sensitivity to luminance contrast across spatial frequencies, improve rapidly over the first postnatal months,^6–11^ constraining the visual information available to the developing cortex. It has been proposed that this transient period of low-resolution input may scaffold later visual development.^12–17^ Individuals who are functionally blind at birth due to dense bilateral congenital cataracts and later undergo sight-restoring surgery rapidly gain access to relatively high-resolution visual input without experiencing the typical progression from initially blurry to higher-resolution vision^18^. These individuals later show characteristic impairments in configural and shape-based processing.^19,20^ Consistent with this account, deep neural networks (DNNs) trained on initially blurred images develop more spatially integrated, shape-based representations.^16,21–23^ Direct empirical evidence that low-resolution early input shapes developing cortical representations, however, remains limited. More fundamentally, the spatial scale of the features contributing to naturalistic object representations in infancy has not been characterized.

Newborns are most sensitive to coarse visual information, carried by low spatial frequencies. By around 9 weeks, however, relative sensitivity across spatial frequencies, captured by the shape of the contrast sensitivity function, approaches that of adults, although absolute contrast sensitivity continues to improve throughout infancy and into adulthood.^9^ Such coarse visual information can support object recognition during infancy. Newborns, for example, can detect global structure, but only when low spatial frequencies are present,^24^ and older infants can categorize objects from silhouettes and shape skeletons.^25,26^ However, these findings do not establish whether infant cortex is tuned to spatially integrated object features. Indeed, mid-level visual functions that require spatial integration, such as segmentation and grouping, remain immature well into the first year.^7^ These functions depend in part on recurrent circuitry, including horizontal and feedback connections,^27,28^ which undergo protracted maturation across infancy.^29–31^ This juxtaposition suggests two candidate developmental trajectories: coarse-scale object-feature tuning could emerge early, scaffolded by the low-resolution input characteristic of early infancy, or it could emerge more gradually as integrative circuitry matures and visual experience accumulates. Whether development follows the same trajectory across the visual hierarchy also remains unknown.

To characterize these developmental trajectories across the visual hierarchy, we took a Developmental NeuroAI approach,^32–34^ combining awake infant fMRI with developmentally inspired DNNs. Awake fMRI has revealed early retinotopic organization and category representations along the infant VVS,^35–41^ but the visual features supporting these representations remain unclear. In the Foundations of Cognition (FOUNDCOG) project,^34,41^ infants viewed naturalistic object images (Figure 1A) during awake fMRI at 2 months, after relatively limited postnatal visual experience, and again at 9 months, following considerable visual maturation and accumulated experience. Importantly, although visual acuity and absolute contrast sensitivity remain immature at 2 months, the stimuli were designed and presented at a large visual angle to support visibility at this age (Figure S1). We have previously shown that object-category representations are already present along the VVS by 2 months and strengthen throughout the first year, and that representations in standard object-trained DNNs align with these infant responses.^41^ These findings established DNN–brain correspondence in infancy and laid the foundation for the present study, in which we use controlled manipulations of model visual experience to probe the development of cortical feature tuning.

**Figure 1.**
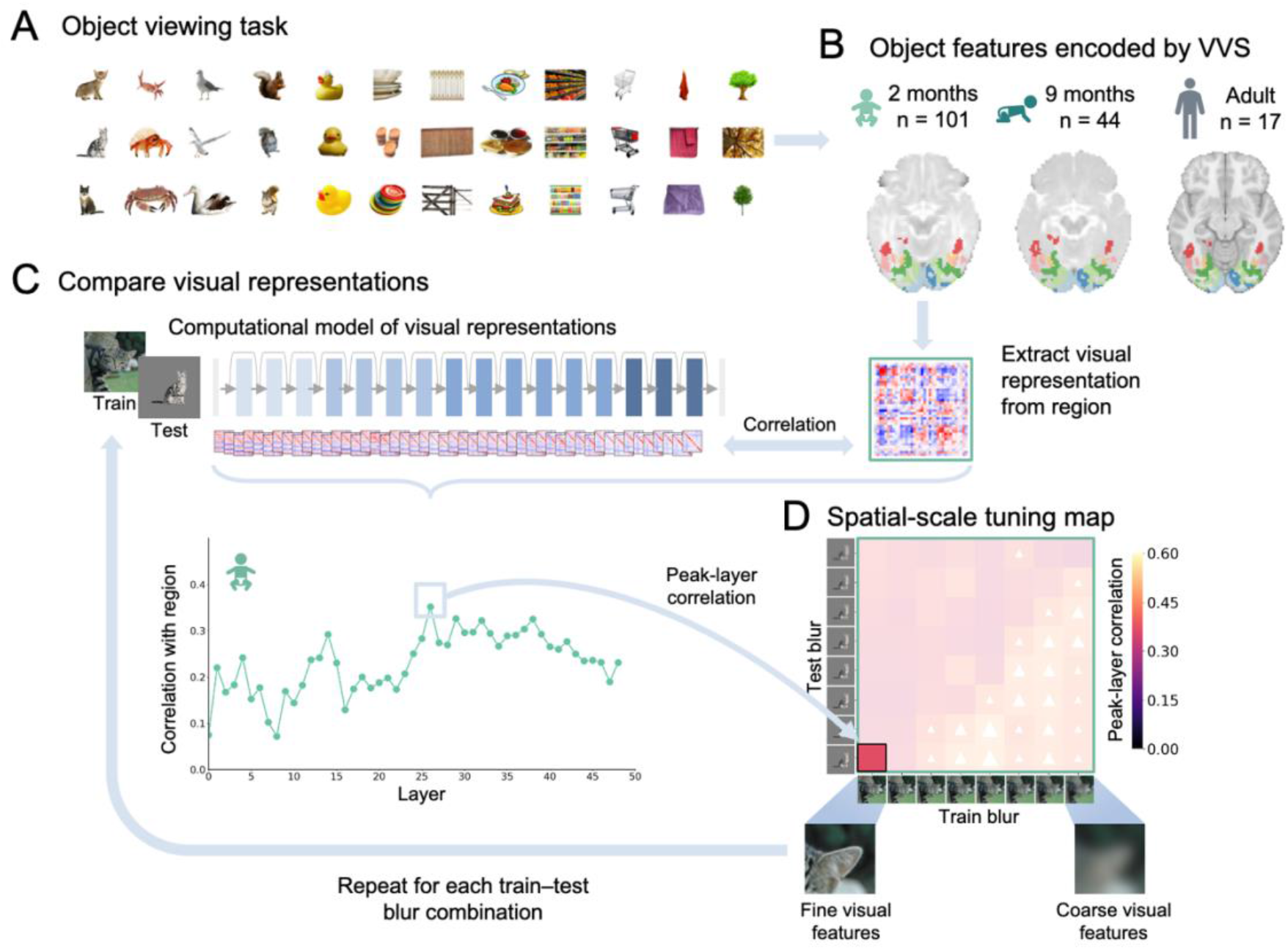
Tuning to coarse and fine object features was characterized along the developing ventral visual stream using awake infant fMRI and blur-trained deep neural networks. Participants viewed 36 images of objects (3 exemplars from 12 categories spanning animate, inanimate large, and inanimate small objects) (A). Functional responses to these stimuli were recorded using fMRI. For each region along the ventral visual stream, group-level visual representations were derived for 2-month-olds (n = 101), 9-month-olds (n = 44), and adults (n = 17) by computing pairwise response similarities between stimuli for each unique combination of runs across participants and averaging across subject-run pairs (B). The same images were presented to a trained deep neural network (DNN; ResNet-50), and layer-wise activation patterns were extracted to generate corresponding model representations. Model–brain correspondence was quantified by comparing the pattern of response similarities in each model layer with that observed for each brain region and age group. The layer yielding the highest correspondence (peak layer) was taken as the model representation that best captured the level of feature complexity represented by a given region (C). To assess tuning to visual features across spatial scales, DNNs were trained and tested on images with varying levels of blur (D). Train blur (x-axis) manipulated the spatial scale of features available during learning, producing models with different learned feature biases, whereas test blur (y-axis) manipulated the spatial scale of visual information available to the trained model at inference. Systematically varying these factors yielded a spatial-scale tuning map for each age group and region.

We exploited a key advantage of DNNs: their learning history and visual input can be systematically manipulated. Standard object-classification training tends to bias models toward local image features,^22,42–44^ whereas training on blurred images restricts fine-scale information and encourages more spatially integrated representations.^16,45–47^ We therefore varied the level of blur during model training (train blur) to manipulate the spatial scale of features available during learning, and independently varied blur when testing the models (test blur) to manipulate the resolution of the current visual input. This allowed learned feature tuning to be dissociated from sensitivity to input resolution (Figure 1). We further compared object-trained models with untrained and retinal-wave- trained networks to ask to what extent early V1 tuning could be accounted for by model architecture or receptive-field properties associated with spontaneous retinal activity.^48–50^ Finally, bandpass-filtered training isolated low-, mid-, and high-spatial-frequency information, allowing us to compare the learning dynamics of artificial visual systems with developmental changes in infant cortex. These complementary manipulations allowed us to characterize how tuning to coarse and fine object features emerges across the developing VVS and to test whether biological and artificial visual systems follow similar developmental trajectories.

## Results

### Blur-trained DNNs reveal spatial-scale tuning in adult visual cortex

DNNs with the ResNet-50 architecture were used for all analyses, as a standard ResNet-50 trained and tested on sharp (unblurred) images showed robust layer-wise correspondence with visual responses across the developing VVS (Figure S4), with peak correlations comparable to or higher than those we previously reported using a supervised AlexNet.^41^ To quantify tuning across spatial scales, we generated a two-dimensional tuning map for each age and visual region from the independently varied train- and test-blur conditions (Figure 1C, 1D). For each train–test blur combination, model–brain correspondence was calculated across DNN layers, and the peak correlation was used to construct the spatial-scale tuning map. As anticipated, models trained on increasingly blurred images exhibited greater sensitivity to lower-spatial-frequency object features (Figure S2), confirming that the training manipulation produced systematic differences in learned feature scale.

We first quantified spatial-scale tuning across the adult VVS, where mature representations provided a benchmark for interpreting developmental differences in infancy. We examined V1 and extrastriate visual cortex (V2–V4), together with later VVS regions VO1 and FG4 (region-of-interest [ROI] definitions are shown in Figure S3). Split-half noise ceilings of the group-level visual representations in each region and age group are reported in Table S1. In adults, responses in V1–V4 were typically best fit by intermediate model layers (Figure 2F and Figure S5) and by models that had learned to extract coarser features through training on intermediate to high levels of blur but were subsequently presented with sharper images at test (Figure 2A). Standard models trained on sharp images, which have access to both coarse- and fine-scale information during training, rely strongly on learned fine-scale local features.^22,42–44^ These models achieved the highest object-classification accuracy (Figure S8A). Despite this, they were consistently among the poorest fits to object responses in these regions. A notable asymmetry emerged in the tuning patterns. Training on blurry images and testing on sharp images produced the strongest correspondence with visual cortex, whereas the reverse combination did not. This extends previous findings that blurry training can increase model alignment with mature brain representations,^46,47^ indicating that coarse learned features and fine visual information at test make complementary contributions to adult object representations across visual cortex.

**Figure 2.**
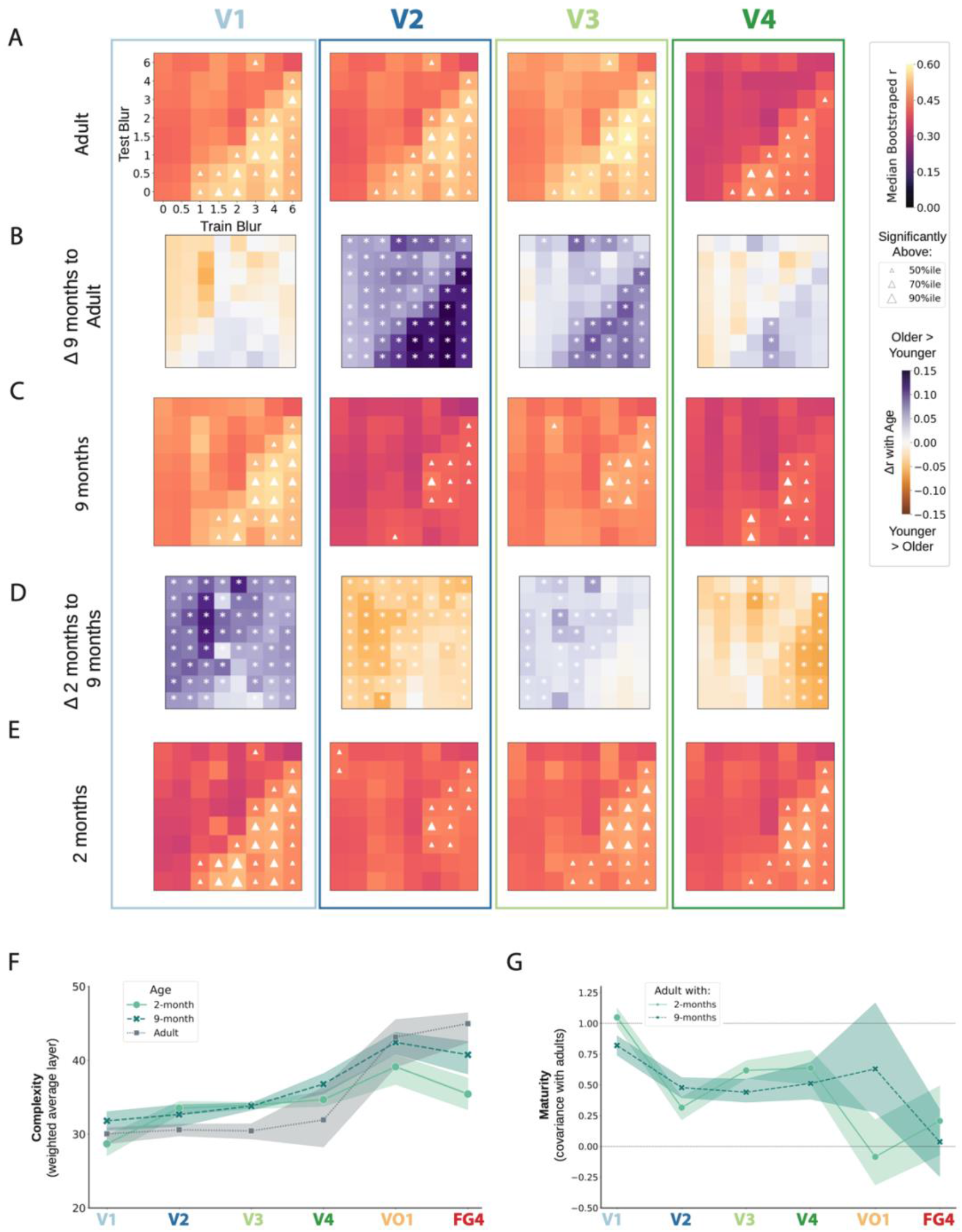
Development of tuning to coarse and fine object features along the ventral visual stream. The pattern of tuning in V1 was already mature by 2 months, whereas tuning in later visual regions showed protracted development through 9 months. Tuning to coarse and fine object features was assessed as described in Figure 1 by training deep neural networks (DNNs) on objects with varying levels of Gaussian blur (train blur, x-axis; σ = 0–6) and testing them across the same range (test blur, y-axis) for adults (A), 9-month-olds (C), and 2-month-olds (E). For each bootstrap and train–test blur combination, model–brain correspondence was calculated across DNN layers, and the layer yielding the highest correspondence was identified as the peak- corresponding layer. Each cell shows the median peak-layer model–brain correlation across 1000 bootstrap resamples of subject-run pairs. Triangles indicate train–test combinations with model–brain correspondence above the 50th, 70th, or 90th percentile across combinations within each tuning map, with larger triangles indicating higher percentiles. Noise-ceiling-scaled tuning maps are shown in Figure S9. Unscaled line plots of the model–brain correlations in (A), (C), and (E), with 95% bootstrap confidence intervals, are shown in Figure S10. Difference maps were calculated by subtracting the younger from the older age group for equivalent train–test blur combinations (B, D); purple indicates greater correspondence at the older age and orange greater correspondence at the younger age. (F) Peak-corresponding layers were summarized across the tuning map by calculating, for each bootstrap, the weighted average of the layer numbers across train–test combinations, weighted by their corresponding model–brain correlations. Medians and 95% bootstrap confidence intervals of these weighted-average layer estimates are shown for each age group. (G) Maturity of tuning was quantified as the covariance between infant and adult tuning maps, scaled by the variance of the corresponding adult tuning map, such that a value of 1 corresponds to the adult reference value. Medians and 95% bootstrap confidence intervals are shown.

Tuning patterns in later VVS regions were distinct from those of early visual cortex and V4 (Figures S6– S7). Consistent with increasingly abstract object representations along the VVS,^51^ responses in VO1 and FG4 were best captured by later model layers (Figure 2F) and by train–test combinations associated with higher object-classification accuracy (Figure S8), rather than by models with access to features at any particular spatial scale. Thus, although our analyses extended into higher-level ventral visual regions, the clearest spatial-scale tuning patterns were observed in V1–V4. Subsequent analyses therefore focused primarily on these regions.

### Infant V1 is tuned to coarse and fine object features by 2 months

Spatial-scale tuning maps were calculated for infants using the same procedure as for adults (Figure 2C, 2E). To quantify maturity relative to adults, we calculated the covariance between infant and adult tuning maps, scaled by the variance of the corresponding adult tuning map. This measure reflects both the similarity of the tuning pattern and its relative amplitude but is insensitive to uniform offsets in model–brain correspondence, with a value of unity corresponding to the adult reference value (Figure 2G). Tuning in V1 was remarkably mature by 2 months, already closely resembling the adult pattern (Figure 2E). Like adults, infant V1 responses were best modeled by intermediate model layers. Scaled covariance between 2-month-old and adult V1 was closely matched to the adult reference value of 1 (median across bootstraps = 1.049, 95% CI = [0.978, 1.125]). Between 2 and 9 months, model–brain correspondence increased broadly across the tuning map rather than selectively for particular train– test blur combinations, while the overall pattern of tuning remained consistent (Figure 2C, 2D). Thus, development between 2 and 9 months primarily reflected a general strengthening of model–brain correspondence rather than a selective change in tuning to features at any particular spatial scale. The adult-like tuning observed in V1, including for sharp test images, suggests that immature visual acuity and contrast sensitivity at 2 months did not prevent fine-scale information available in our stimuli from contributing to V1 object representations. This reflects, at least in part, the stimulus design. Objects were presented at a wide visual angle to support visibility at 2 months and reduce the influence of age-related test-time visual limitations on the recorded object responses.

Given the mature V1 tuning observed by 2 months, we asked whether greater postnatal visual experience was associated with a different tuning pattern. We therefore separately examined a subgroup of FOUNDCOG participants who were born preterm (n = 17). These infants were scanned at a similar corrected age to term-born controls but had approximately 8 additional weeks of postnatal visual experience (Figure S11A, S11B). All infants in the preterm group had spent time in the neonatal intensive care unit (NICU) after birth, so this comparison cannot isolate the effect of visual experience alone. Nevertheless, V1 tuning was highly similar between groups (Figure S11C), providing no evidence that greater cumulative postnatal visual experience was associated with a corresponding difference in V1 tuning at 2 months corrected age.

### Retinal-wave-derived features alone do not account for object-feature tuning in infant V1

The mature spatial-scale tuning observed in V1 by 2 months raised the question of its developmental origins. It may partly reflect early circuit organization established through genetic constraints and spontaneous neural activity, potentially alongside visual experience before birth.^52^ Retinal waves contribute to the establishment of early V1 receptive-field properties and, in animal models, play a role in establishing retinotopy and orientation selectivity prior to eye opening^48,49^. DNNs trained on classes of retinal waves have similarly been shown to exhibit V1-like receptive-field morphology.^50,53^ If receptive-field properties established through spontaneous retinal activity were sufficient to account for infant V1 responses, a retinal-wave-trained DNN should capture these responses as well as an object-trained network. Alternatively, retinal-wave-driven organization may provide an initial scaffold, with prenatal visual input and subsequent postnatal experience with objects contributing additional features. To test these possibilities, we compared infant visual responses with previously published DNNs trained on simulated retinal waves (Figure 3A),^50,54^ trained only on standard, sharp object images, or pretrained on retinal waves followed by fine-tuning on the same sharp object images. Responses were also compared with an untrained network to control for effects of model architecture.

**Figure 3.**
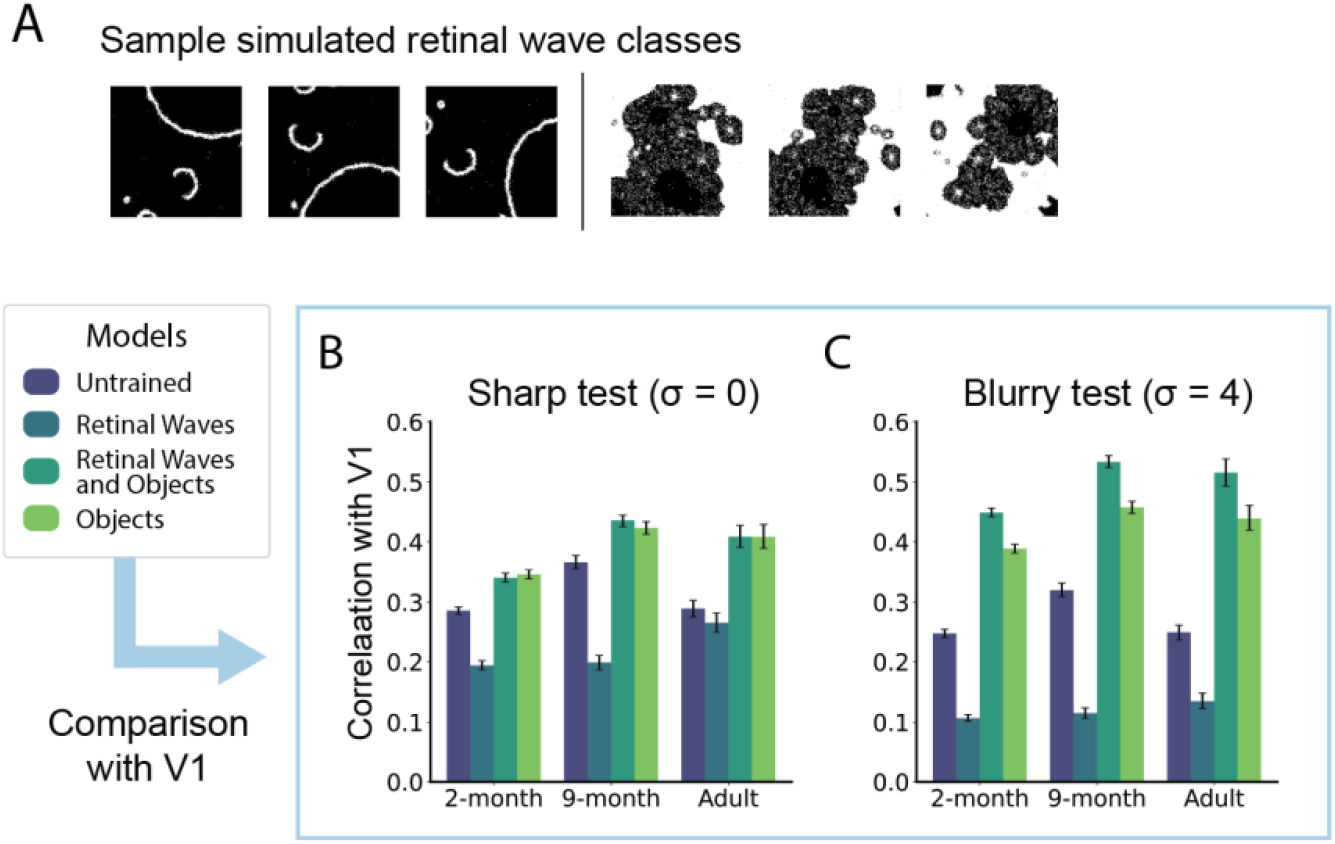
**Retinal-wave training alone does not account for object-feature tuning in infant V1**. To test whether early V1 representations could be explained by features learned from retinal waves, object responses were compared with deep neural networks (DNNs) that were trained on simulated retinal waves (blue), trained on objects (light green), pretrained on retinal waves and subsequently fine-tuned on objects (dark green), or left untrained (purple). The trained models were developed and publicly released by Cappell and colleagues.^54^ Example retinal-wave stimuli are shown from the Retinal Wave Dataset^55^ (CC BY 4.0; resized for visualization) (A). For each model, age group, and test-blur condition, layer-wise model–brain correspondence was calculated for V1 using Spearman correlation across 1000 bootstrap resamples per cohort. The layer yielding the highest correspondence was identified as the peak-corresponding layer, and the corresponding model–brain correlation was retained for comparison. Sharp (σ = 0; B) and blurry (σ = 4; C) test stimuli are shown here as illustrative conditions. V1 results for all test-blur levels are shown in Figure S14A. Plotted values show the bootstrap median peak-layer model– brain correlation, with error bars indicating 95% bootstrap confidence intervals. Corresponding results across all ventral visual stream regions and test-blur levels, together with median peak- corresponding layer estimates, are shown in Figure S15.

The retinal-wave-trained model was a significantly poorer model of V1 than the object-trained model across all ages when tested on either sharp or blurry stimuli (Figure 3B, 3C; see Figure S14A for all test- blur levels), indicating that retinal-wave-like training alone was insufficient to account for V1 tuning, even at 2 months. The same patterns were observed with grayscale FOUNDCOG stimuli (Figure S14B), indicating that the advantage of object-trained models was not explained by access to color information. The untrained network was likewise a poorer model of V1 than the object-trained model (Figure 3B, 3C), indicating that model architecture alone could not account for V1 tuning. For sharp test stimuli, retinal-wave pretraining conferred no benefit over object training alone (Figure 3B). For blurry test stimuli, however, the model pretrained on retinal waves and subsequently fine-tuned on objects showed greater model–brain correspondence than the object-trained model for both infants and adults (Figures 3C and S14A).

### Extrastriate visual cortex shows protracted development of spatial-scale tuning

In contrast to V1, which exhibited a mature tuning pattern by 2 months, spatial-scale tuning in V2–V4 remained distinct from adults through 9 months, indicating protracted development in extrastriate visual cortex (Figure 2A–E, 2G). Across V2–V4, infant responses tended to be best captured by models trained with greater levels of blur, with relatively weaker correspondence for sharp test conditions than was evident in the adult tuning patterns. This pattern was clearest in V2 and V3, where model– brain correspondence for the train–test blur combinations that best captured adult visual representations was systematically lower in both 2- and 9-month-old infants (Figure 2A, 2C, 2E). Within the infant tuning maps, models trained with high levels of blur and tested at intermediate blur levels tended to provide the best fits. V4 followed a somewhat different developmental pattern. At 2 months, models trained with the highest level of blur were among the best-fitting models, whereas by 9 months correspondence with these models was reduced and the adult-like tuning pattern had begun to emerge. However, this apparent spatial-scale bias was not consistently evident in the age-difference maps (Figure 2B, 2D). We therefore tested the apparent contribution of coarser-scale features more directly using spatial-frequency-specific, bandpass-filtered models below. As in V1, intermediate model layers tended to provide the best fits to V2–V4 across age groups (Figure 2F). These developmental differences could not readily be explained by differences in motion between cohorts, as the same qualitative pattern was observed in a low-motion subset of infant runs (Figure S12).

In later VVS regions, VO1 and FG4, development was characterized by the progressive emergence of the adult pattern of model correspondence. At 2 months, responses showed relatively little differentiation across train–test blur combinations and were captured by earlier model layers, particularly in FG4, consistent with representations of lower feature complexity at this age and our previous finding that category structure in ventral temporal cortex strengthens progressively from infancy to adulthood.^41^ By 9 months, tuning had become more differentiated and adult-like, particularly in VO1, with increased correspondence for later model layers and train–test combinations associated with higher classification accuracy (Figures 2F, S6, and S8). A similar pattern was observed in the low-motion subset of infant runs (Figure S13).

### Low-spatial-frequency representations take longest to converge with visual cortex during learning

In the blur-trained models, the strongest correspondence with visual cortex (V1–V4) was observed when networks were trained on blurry images but tested on sharp images. This asymmetry suggested that representations based on lower spatial frequencies may require more learning, whereas higher- spatial-frequency information can contribute strongly at test despite being attenuated during training. We therefore hypothesized that representations derived from low-spatial-frequency information would take longer to converge with visual cortical responses, whereas higher-spatial-frequency correspondence would emerge earlier and might be present, even before object learning, in the untrained model. To test this directly, we trained separate DNNs on bandpass-filtered ImageNet images containing low-, mid-, or high-spatial-frequency information (Figure 4A). A preliminary analysis across 60 epochs of training showed that model–brain convergence occurred very early, with the midpoint of learning typically falling within the first epoch (Figure S16), consistent with previous work showing that model–brain correspondence emerges rapidly during DNN training.^56^ We therefore defined the first three epochs as the critical training window, encompassing the period of rapid representational change while allowing these early learning dynamics to be sampled at high temporal resolution. Model representations were densely sampled throughout this window and compared with developing visual responses (Figures 4B and S17).

**Figure 4.**
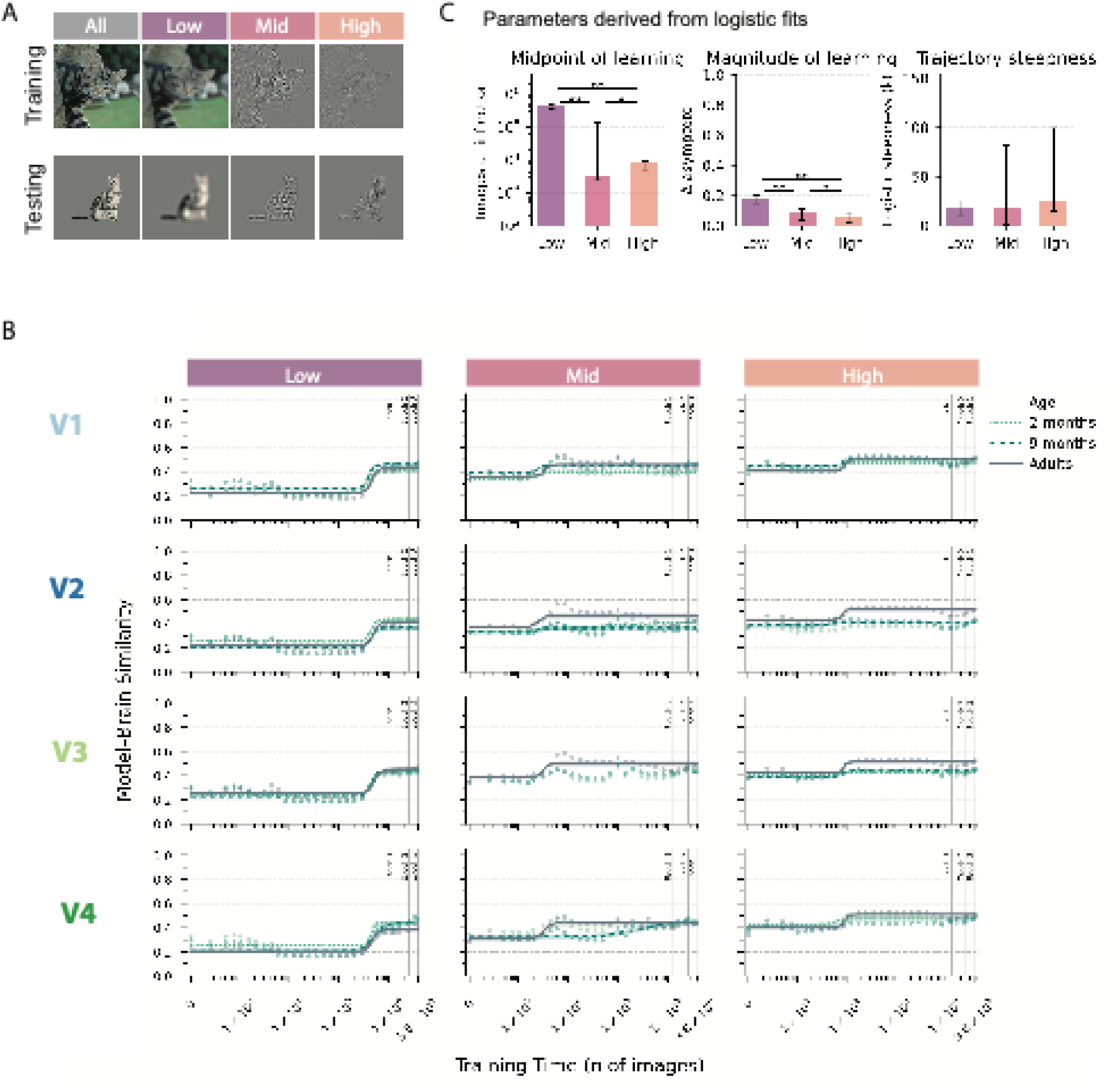
Low-spatial-frequency representations take longest to converge with visual cortex during object learning. (A) Example ImageNet training image and FOUNDCOG test stimulus decomposed into low-, mid-, and high-spatial-frequency bands. (B) Logistic fits of model–brain correspondence across the critical training window. At each sampled training checkpoint, model representations were compared with responses from V1, V2, V3, and V4 in 2-month-olds, 9-month-olds, and adults. For each age by ROI by spatial-frequency-band combination and bootstrap sample, model–brain correlations were averaged across the 10 independently initialized network instances within each layer before selecting the peak-corresponding layer. Peak-layer model–brain correspondence was normalized by the age- and ROI-specific noise ceiling. Points show the bootstrap median at each sampled checkpoint. Logistic functions were fitted separately to each bootstrap training trajectory. For combinations meeting the prespecified fit-reliability criterion (≥80% usable bootstrap fits; 31/36; Figure S19B), the median logistic fit and 95% bootstrap interval are shown. (C) Summary parameters were derived from the reliable logistic fits: midpoint of learning, magnitude of learning, and trajectory steepness. Bars show the median and 2.5th to 97.5th percentile range across reliable age by ROI combinations. Asterisks indicate significant pairwise differences between bands (Holm-corrected Wilcoxon signed-rank tests; Table S4). Age- and ROI-specific estimates are shown in Figure S20. Low-spatial-frequency representations reached their midpoint of learning latest and showed the greatest magnitude of learning, whereas mid- and high-spatial-frequency representations reached their midpoint substantially earlier. Trajectory steepness did not differ significantly across bands.

As hypothesized, the untrained model already showed significantly higher peak-layer model–brain correspondence for the high band than for the mid or low bands across all visual cortex regions (V1– V4) and age groups (Figure S18; Table S2). Model–brain correspondence did, however, increase significantly over the critical training window for every age-by-ROI-by-band combination (Table S3), demonstrating a contribution of learning across all three spatial-frequency bands. Logistic functions were fitted to these trajectories, from which the midpoint of learning, magnitude of learning, and trajectory steepness were derived (Figure 4B, 4C). Reliable fits were obtained for 31 of 36 age-by-ROI-by-band combinations (Figures 4B and S19). Parameter estimates summarized across age and ROI are shown in Figure 4C (Table S4). Age-by-ROI-specific estimates are shown in Figure S20, with corresponding pairwise differences across spatial-frequency bands reported in Table S5.

Across reliable trajectories, the midpoint of learning generally occurred within the first epoch but differed markedly across spatial-frequency bands. The mid band reached its midpoint earliest (median = 3.2 × 10^3^ images), followed by the high band (median = 8.3 × 10^3^ images), whereas the low band reached its midpoint substantially later (median = 4.4 × 10^5^ images), approximately 1.7 orders of magnitude later than the high band (Figure 4B, 4C; Table S4). The magnitude of learning also differed across bands, with the low band showing the greatest increase, the high band the smallest, and the mid band an intermediate increase. Trajectory steepness did not differ significantly across bands. Thus, although learning contributed to representations across all three spatial-frequency bands, high-spatial-frequency representations corresponded strongly with visual cortex from early in training, whereas low-spatial-frequency representations changed most with learning and converged last.

### Infant extrastriate cortex is biased toward coarser features relative to adults

Having established distinct learning dynamics across spatial-frequency bands, we next asked whether low-, mid-, and high-spatial-frequency representations corresponded differently with infant and adult visual responses at the end of the critical training window (Figure 5A; Table S6). Low-band correspondence peaked at a consistent depth across ages, shifting from intermediate layers in V1–V3 to late layers in V4, in line with V4’s higher position in the visual hierarchy and consistent with more spatially integrated features. Mid- and high-band correspondence peaked in the earliest model layers in V1 at all ages and in adult V2–V4. In infants, however, mid- and high-band peaks occurred in later layers in V3 and V4 at both ages and in V2 at 2 months (Figures S17, S21). Direct comparison of the layer-wise model representations further indicated that the peak-corresponding low- and high-spatial- frequency representations contained distinct representational structure and therefore captured non- redundant information (Figure S22).

**Figure 5.**
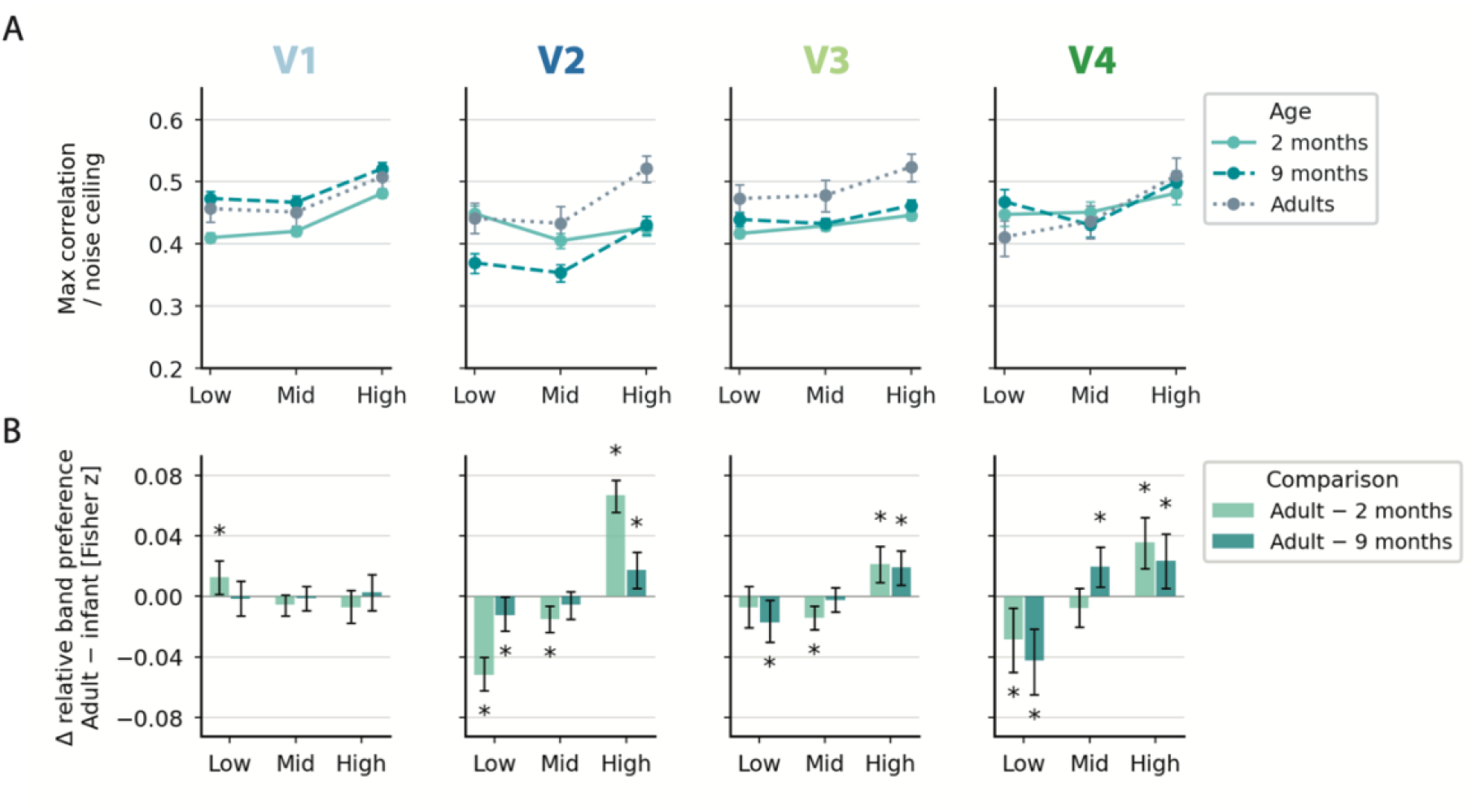
Infant extrastriate cortex shows a relative low-spatial-frequency bias through 9 months. (A) Band-specific peak-layer model–brain correspondence at the end of the critical training window across V1, V2, V3, and V4 in 2-month-olds, 9-month-olds, and adults. Model–brain correspondence was normalized by the age- and ROI-specific noise ceiling. Points show bootstrap medians and error bars show 95% bootstrap confidence intervals. Peak- corresponding layers are shown in Figure S21. (B) Developmental differences in relative spatial-frequency-band preference at the end of the critical training window. Fisher-transformed model–brain correlations were demeaned across bands within each age group before comparing infants with adults, thereby isolating relative band preference from overall differences in model–brain correspondence across ages. Positive values indicate relatively greater correspondence in adults and negative values relatively greater correspondence in infants. Error bars show 95% bootstrap confidence intervals, and asterisks indicate intervals excluding zero. In V2–V4, adults showed relatively greater correspondence with high-spatial-frequency representations than infants at either age, whereas infants showed relatively greater correspondence with low-spatial-frequency representations in five of six comparisons. This relative low-spatial-frequency bias remained evident at 9 months.

To isolate relative band preference from overall age differences in model–brain correspondence, Fisher-transformed correlations were demeaned across bands within each age group before comparing infants with adults. In V1, relative band preference was close to adult values at both ages, the only difference being a small reduction in relative low-band correspondence at 2 months (Figure 5B; Table S7). In V2–V4, by contrast, adults showed relatively greater correspondence with high-spatial-frequency representations than infants at either age, whereas infants showed relatively greater correspondence with low-spatial-frequency representations in five of six comparisons. This relative low-spatial-frequency bias remained evident at 9 months. Biological and artificial visual development therefore diverged: low-spatial-frequency representations, the last to converge with visual cortex during model training, corresponded relatively more strongly with infant than adult extrastriate cortex at both 2 and 9 months.

## Discussion

To characterize the development of tuning to coarse and fine object features in infant visual cortex, we used a Developmental NeuroAI approach combining awake infant fMRI with developmentally inspired DNNs. This revealed distinct developmental trajectories across the visual hierarchy. By 2 months, V1 already exhibited a mature tuning pattern sensitive to both coarse and fine object features, whereas tuning in extrastriate visual cortex (V2–V4) remained distinct from adults through 9 months. Critically, this protracted development reflected a relative bias toward coarse, low-spatial-frequency features that was not readily explained by developing acuity. This bias also revealed a striking divergence from object-trained DNNs, in which representations derived from low spatial frequencies, those relatively emphasized in infant extrastriate cortex, took longest to converge with visual cortex during training.

Notably, neither candidate developmental trajectory fully captured the observed pattern. Coarse-scale object information was already robustly represented by 2 months, with a relative coarse-feature bias evident in extrastriate cortex despite the protracted maturation of horizontal and feedback connections,^29–31^ indicating that mature integrative circuitry is not required for coarse-scale tuning to emerge. Conversely, the mature multi-scale tuning already evident in V1 suggests that the extrastriate coarse-feature bias is not simply inherited from limitations in acuity or contrast sensitivity. Our findings cannot, however, determine whether low-resolution visual experience early in development contributes to establishing the coarse-scale tuning observed in infancy.^16,57^ Rather, they show that coarse- and fine-feature tuning is shaped by the cortical hierarchy, following distinct developmental trajectories in V1 and extrastriate cortex.

The mature V1 tuning observed by 2 months closely resembled adult sensitivity to both coarse and fine features. Although the magnitude of model–brain correspondence strengthened between 2 and 9 months, the pattern of tuning remained stable, suggesting that subsequent development primarily strengthened an already established representational structure rather than substantially altering its spatial-scale tuning. This strengthening coincides with a period of rapid synaptogenesis in human V1 during infancy.^58^ The maturity of V1 tuning, including sensitivity to fine-scale information available in sharp test images, also provides important internal validation that the spatial-frequency information probed by our models could contribute to cortical responses even at the youngest age tested. Importantly, untrained and retinal-wave-trained models showed substantially weaker correspondence with infant V1 than object-trained models, indicating that neither model architecture alone nor features learned from spontaneous retinal activity were sufficient to account for the mature tuning observed by 2 months. Retinal-wave pretraining did, however, selectively enhance correspondence for blurry test stimuli, suggesting that structured spontaneous activity may shape representations in ways that subsequently interact with visual experience.

At the time of their first scan, infants had already been exposed to the natural statistics of the visual world for more than two months. The greater correspondence with object-trained models could therefore indicate that the structure of postnatal visual input over the first two months contributes to shaping V1 responses. Electrophysiological evidence from kittens has shown that V1 receptive fields are sensitive to visual input statistics and are fine-tuned according to the early visual environment.^59^ Similarly, in humans, visual experience during the early postnatal months contributes to the establishment of fine-scale retinotopic organization in early visual cortex, including V1.^60^ Alternatively, correspondence with object-trained DNNs need not imply that the biological tuning they capture was itself learned through visual experience. DNNs can capture both innate and experience-dependent properties of neural representations.^61,62^ The object-trained models may therefore capture components of V1 tuning established prior to or independently of postnatal visual experience that are not accounted for by the retinal-wave-trained model. Notably, the pattern of V1 tuning at 2 months was similar across term-born controls and preterm-born infants who had spent time in the NICU, despite the latter group having substantially more postnatal visual experience. Future work is required to disentangle how prenatal constraints and early visual experience interact to establish mature object-feature tuning in V1 so early in development.

In contrast to V1, infant extrastriate cortex exhibited protracted development, remaining distinct from adults through 9 months. Bandpass analyses revealed that this immaturity reflected a relative bias toward coarser, low-spatial-frequency features that persisted through 9 months. Importantly, this was a relative bias rather than an absence of sensitivity to finer-scale information. The bias cannot simply be explained by immature acuity, as the spatial frequencies in our stimuli (up to approximately 3.2– 4.9 cycles per degree [cpd]) lie within VEP-based acuity estimates by 2 months and are readily resolvable by 9 months.^8,10^ The dissociation between V1 and extrastriate cortex further argues against an origin earlier in the visual pathway. If the relative coarse bias arose in the optics, retina, lateral geniculate nucleus or V1 itself, it should be evident in V1 responses. Instead, V1 showed adult-like tuning across spatial scales by 2 months. Although we cannot exclude a contribution from residual differences in contrast sensitivity, which remains lower in infants than in adults^9^, such a contribution would be expected to affect V1 as well as extrastriate cortex. Similar noise ceilings across ages in V1– V4 further argue against lower reliability of infant responses as an explanation. Related EEG work in 6–8-month-old infants similarly found greater correspondence with lower-spatial-frequency information, which was interpreted as consistent with developmental differences in spatial sensitivity.^63^ Using fMRI, we localize such a bias within the visual hierarchy, showing that it is present in extrastriate cortex but not in V1, where tuning across spatial scales is already mature.

A sustained coarse-feature bias may be beneficial for early visual learning. Coarse-to-fine perceptual development has been proposed to provide a functional inductive bias supporting the development of spatial integration and shape-based representations,^12,14,16,21,23^ while coarser features can facilitate category learning by increasing within-category similarity and accentuating between-category differences.^14,21^ Our findings suggest that such a bias is maintained within extrastriate cortex even as sensitivity to finer visual information increases. This raises the hypothesis that the prolonged coarse-feature bias supports category learning across the first year by shaping the visual information propagated to later object-selective regions as category representations continue to develop.^41^

The learning dynamics of object-trained DNNs showed the opposite bias to that of developing extrastriate cortex. Representations derived from high-spatial-frequency information corresponded strongly with visual cortex from early in training, whereas low-spatial-frequency representations changed most with learning and converged last. High-spatial-frequency correspondence was evident even in the untrained network, where it exceeded that of the mid and low bands across regions and ages, and it changed least with learning, consistent with a contribution from the model architecture rather than learned object features alone. These frequency-specific dynamics also help explain why models trained on blurry images but tested on sharp images showed the strongest correspondence with visual cortex. Blurry training promoted the learning of coarser object features, while fine visual information available at test made an additional contribution. Previous work has implicated both training regime and architecture in the local-feature bias of standard convolutional neural networks.^42,64–66^ Our findings suggest that both contribute. Visual diet altered the spatial scale of learned features, but higher-spatial-frequency representations nevertheless showed greater correspondence with cortical responses than lower-frequency representations, even without learning.

The distinct developmental trajectories across V1 and extrastriate cortex also have implications for how visual development is modelled. DNNs typically optimize the visual hierarchy concurrently, whereas biological development may involve regionally distinct periods of maturation and plasticity. Modeling visual development as a uniform optimization process across the hierarchy may therefore miss an important feature of cortical development. Incorporating staged learning dynamics, for example through layer-specific learning schedules or sequential critical periods, may provide a more developmentally informed approach. Sequential critical periods have already been shown to promote more stable representation learning in DNNs.^67^

There are several important limitations when interpreting these findings. First, the stimuli were designed and presented at a large visual angle to support visibility at 2 months and contained a maximum spatial frequency of 4.9 cpd. Our conclusions therefore concern spatial-frequency information available at this age and cannot address tuning to finer spatial details that become available later in development. Second, participants viewed only the original broadband stimuli. Blurred and bandpass-filtered images were presented only to the computational models. We therefore inferred the contribution of different spatial scales from model–brain correspondence rather than measuring cortical responses to filtered stimuli directly. An adult fMRI study using broadband, low-pass, and high-pass filtered objects similarly reported stronger responses in V1–V4 to high- than low-spatial-frequency objects,^68^ consistent with the strong contribution of fine-scale information to adult V1–V4 representations observed here. Direct manipulation of spatial-frequency content in infants would be needed to establish whether comparable developmental differences are observed when frequency content itself is varied experimentally.

Third, our analyses characterized shared representations at the group level within atlas-defined ROIs. These ROIs were based on adult cytoarchitectonic parcellations normalized to age-appropriate infant templates, limiting localization precision, particularly across neighboring V2–V4. Any contamination by the more mature V1 representation would, however, tend to make infant extrastriate tuning appear more adult-like and therefore work against the observed developmental differences. Group- and ROI-level averaging improved stability but obscured individual differences and within-region heterogeneity. The preterm comparison should also be interpreted cautiously. With only 17 infants, the absence of a detectable difference provides limited evidence about the effect of additional postnatal visual experience.

Fourth, our computational analyses were based primarily on ResNet-50 models trained on ImageNet. Whether the developmental patterns observed here generalize across model architectures and training datasets remains to be established. The effects of blurry training have been shown to generalize across convolutional and transformer architectures,^23,47^ suggesting that this manipulation is not specific to ResNet-50. Finally, comparisons between learning trajectories in DNNs and development in infants should be interpreted with caution. Learning in DNNs was indexed by the number of training images viewed, whereas infant development was measured in months. Although approaches have been proposed for relating model training exposure to developmental time,^69,70^ the correspondence between the two is not straightforward, given differences in the nature, diversity, and temporal structure of experience in infants and DNNs. Our comparison therefore concerns the relative order and dynamics with which coarse- and fine-scale representations emerge, rather than a direct correspondence between model training time and infant age.

Relatedly, the models used here were trained on fixed image transformations rather than developmental trajectories. This provided a tractable way to isolate the contribution of features at different spatial scales while avoiding potential continual-learning effects as higher-spatial-frequency information becomes increasingly available.^22^ A next step will be to combine the region-specific developmental biases identified here with more realistic trajectories of visual experience. Infant visual input changes along multiple dimensions, including perceptual fidelity and semantic content, whose developmental trajectories may form curricula that support early learning.^12,14,16,70–75^ Developmentally informed models could extend this trajectory back into the prenatal period, incorporating spontaneous retinal activity and structured sensory input such as light exposure in utero,^52^ before progressing through initially degraded postnatal input to increasingly rich visual experience. Combined with awake infant fMRI, such models could provide a framework for testing how proposed developmental visual curricula shape emerging representations.

This work revealed a hierarchical developmental trajectory of object-feature tuning across spatial scales in infant visual cortex, with mature tuning in V1 by 2 months alongside a coarse-feature bias in extrastriate cortex that persisted through 9 months. The divergence between this sustained biological bias and the rapid emergence of higher-spatial-frequency correspondence in object-trained DNNs highlights distinct inductive biases in biological and artificial visual development. More broadly, this work illustrates the potential of developmentally inspired DNNs to reveal the features encoded by the developing ventral visual stream and to probe the mechanisms of early visual learning.

## Acknowledgments

This project has received funding from the European Research Council (ERC) under the European Union’s Horizon 2020 research and innovation programme (grant agreement No. 787981, FOUNDCOG; R.C.). Support was also provided by Taighde Éireann – Research Ireland Frontiers for the Future grant 22/FFP-A/11050, Freeze Motion (R.C.), Taighde Éireann – Research Ireland (formerly Irish Research Council) Government of Ireland Postgraduate Scholarship grants GOIPG/2023/2479 (Á.T.D.) and GOIPG/2021/223 (C.O’D.), and the Science Foundation Ireland grant RC-17/RCPhD/3482-RCPhD (A.K.). Thank you to Chiara Caldinelli, Tamrin Holloway and Jessica White for their significant contributions to data collection, and to Lorijn Zaadnoordijk for her contribution to conceptualization of the FOUNDCOG paradigm and acquiring FOUNDCOG ethical approval. We would like to thank Sojo Joseph, radiographer at the Trinity College Institute of Neuroscience for his contribution to infant and adult MRI acquisition and the extended infant data collection team: Claire Ambre, Katie Herbert, Josué Rico Picó, and Anisha Wadhwa for their assistance with acquisition, and Michelle Bergin, Alessandra Biagini, Lucy Corcoran, Niamh Dunne, Linda Farren, Siobhán Lee and Deirdre O’Hara, for clinical supervision during infant scans. Thank you to the members of the Cusack Lab who provided feedback on earlier versions of this work, in particular Marie Santillo, Kevin Campion and Cameron Murphy, and to Calum Kyle for his contribution to literature review. We wish to sincerely thank all of our FOUNDCOG participants and their families who so generously gave of their time to support this project, without whom this research would not have been possible.

## Author contributions

Conceptualization: Á.T.D., C.O’D., A.Tr., A.K., R.C.; methodology: Á.T.D., C.O’D., A.Tr., A.K., R.C.; resources: Á.T.D., A.Tr., G.K., E.-L.D., A.Ta., A.T.B., A.F., E.J.M., R.C.; investigation: Á.T.D., C.O’D., A.Tr., A.K., G.K., E.-L.D., R.C.; project administration: Á.T.D., A.Tr., G.K., E.-L.D., A.Ta., A.T.B., A.F., E.J.M., R.C.; data curation and software: Á.T.D., C.O’D., A.Tr., G.K., R.C.; formal analysis and validation: Á.T.D., R.C.; writing – original draft: Á.T.D., R.C.; writing – review & editing: Á.T.D., C.O’D., A.Tr., A.R.W., R.C.; visualization: Á.T.D., C.O’D., R.C.; supervision: E.J.M., R.C.; funding acquisition: R.C.

## Supplemental information

**Document S1**. Supplemental methods, Figures S1–S22, Tables S1–S7.

## Declaration of interests

The authors declare no competing interests.

## Declaration of generative AI and AI-assisted technologies

The authors wrote the text and developed the code as described in the Author contributions. During the preparation of this work, the authors used ChatGPT (OpenAI) and Claude (Anthropic) for editing sections of the text and formatting code scripts. The authors reviewed and edited the output as needed and take full responsibility for the content of the published article.

Methods Resource availability Lead contact

Further information and requests for resources should be directed to the lead contact, Áine T. Dineen.

## Materials availability

This study did not generate new materials other than data and code.

## Data and code availability

- Model weights and model–brain correlation values have been deposited at Zenodo (doi:10.5281/zenodo.23326150) and are publicly available as of the date of publication. This paper analyzes infant and adult fMRI data from the FOUNDCOG project, described previously.^41^ Data from infants whose caregivers consented to public data sharing (n = 78 2- month-olds; n = 34 9-month-olds) are available at OpenNeuro (ds006883; doi:10.18112/openneuro.ds006883.v1.0.0). In accordance with our ethical approval, individual-participant fMRI data from the preterm (NICU) group cannot be deposited in a public repository. The group-averaged representational similarity matrix (RSM) for this group is included in the Zenodo record.
- All original code has been deposited at GitHub (https://github.com/dineenai/foundcog_coarse_fine) and is publicly available as of the date of publication.
- Any additional information required to reanalyze the data reported in this paper is available from the lead contact upon request.

### Participants

This study was conducted as part of the Foundations of Cognition (FOUNDCOG) project, a longitudinal neurodevelopment study including MRI scans at approximately 2 and 9 months corrected gestational age (CGA), referred to as the 2-month and 9-month timepoints respectively. Ethical approval was granted by the Trinity College Dublin School of Psychology Research Ethics Committee (SPREC012020- 10), and the ethics committees of the two collaborating maternity hospitals: the Research Ethics Committee of The Rotunda Hospital (REC-2019-018) and the Research Ethics Committee of The Coombe Women and Infants University Hospital (Study No. 8 – 2019). Participants were recruited from The Coombe Hospital and the Rotunda Hospital into one of two cohorts: the term-born control group or the preterm-born neonatal intensive care unit (NICU) group. To be eligible for either cohort, infants were required to have no known congenital abnormalities, visual impairments, or contraindications to MRI. Infants in the control group were born at term (37–42 weeks of gestation) and had no diagnosed brain injury. Infants in the preterm group were born preterm and had spent time in the NICU. These infants were screened for retinopathy of prematurity (ROP) and were excluded if ROP reached Stage 3 or required laser or injection treatment. Eligible infants were invited to attend MRI sessions at approximately 2 and 9 months CGA. Further details of recruitment and study implementation are provided elsewhere.^34^ Informed consent was obtained from a caregiver on behalf of each infant.

Unless otherwise stated, control and preterm cohorts were combined for analysis. Of the 134 infants (29 preterm) who attended the 2-month scan with their caregivers, awake functional MRI (fMRI) data were successfully acquired from 131, who were subsequently invited to return at 9 months. Of these, 104 returned (79%), with 27 (21%) lost to follow-up. Awake fMRI data were successfully acquired from 66 infants at 9 months (63%). Two visual tasks were conducted during awake scanning: a videos task and a pictures task which comprised a series of images of objects. The analyses presented in this manuscript used responses acquired during the pictures task. A subset of 113 2-month-olds (median CGA = 2.33 months, IQR = [2.13, 2.67]; 44 female) and 51 9-month-olds (median CGA = 9.20 months, IQR = [8.95, 9.45]; 23 female) participated in the pictures task. Following preprocessing and motion thresholding, a final sample of 101 2-month-olds (median CGA = 2.33 months, IQR = [2.13, 2.63]; 37 female) and 44 9-month-olds (median CGA = 9.20 months, IQR = [8.92, 9.51]; 22 female), in addition to 17 adults, provided data that were used to calculate group-level visual object representations analyzed here and in related work.^41^

### Recording object responses with fMRI

Whole-brain fMRI responses were recorded while participants viewed 36 objects comprising three exemplars from 12 categories spanning animate, inanimate large, and inanimate small classifications (Figure 1A). Objects were presented in a pseudo-random order, such that one exemplar from each category was presented prior to another exemplar from the same category. Each object was presented for 3 s, interleaved with a fixation cross for 0.5–1.5 s. Each run contained two repetitions of the objects. An attention getter preceded each repetition. Infants contributed a median of 1.5 runs of the pictures task at 2 months and 1 run at 9 months.^34^ Details of the scanning setup and acquisition parameters are provided in the Supplemental Methods.

Objects were segmented, displayed on a perceptually transparent background and projected directly onto the bore of the scanner above the infant’s eyes. Objects loomed toward the infant to hold their attention, initially appearing in the center of the display at half of their maximum height and increasing monotonically in size according to a logarithmic function, such that they spent most of each presentation near their maximum size. The maximum object size was defined as the size at which either its width or height matched the corresponding dimension of the display (29 cm × 16.3 cm; approximately 56° × 34° of visual angle at a viewing distance of approximately 27 cm). Stimuli were rendered at 640 × 360 px, giving a theoretical maximum spatial frequency of 5.7 cpd. However, stimuli were geometrically pre-distorted to correct for the projection angle and bore curvature (Supplemental Methods), so objects did not fill the full 640 × 360 px frame. Along the dimension that matched the display at maximum size, objects spanned 216–550 px (median 227 px), corresponding to a maximum presented spatial frequency of approximately 3.2–4.9 cpd (median 3.4 cpd). Given these display parameters, the stimuli were designed to support visibility at 2 months. An illustrative simulation of the estimated appearance of a sample FOUNDCOG object at this age is shown in Figure S1B. The FOUNDCOG experimental paradigm is available at https://github.com/ClionaOD/foundcog_paradigm.

### Blur-trained deep neural network models

#### Model training

ResNet-50^76^ models were trained on the ImageNet-1K dataset^77^ using supervised object classification. Models were trained on visual diets consisting of images blurred with a fixed level of Gaussian blur, ranging from no blur (sharp) to strongly blurred (σ = 0, 0.5, 1, 1.5, 2, 3, 4, 6 px). Blur was applied after images were randomly cropped to 224 × 224 px. Models were trained for 60 epochs using a standard ImageNet training protocol. Two independently initialized models were trained for each level of blur to control for idiosyncratic variability in the representations learned by individual model instances.^78^

#### Characterizing the spatial frequency of visual features learned by blur-trained DNNs

To verify that training blur altered the spatial scale of features learned by the models, we characterized the spatial-frequency content of image patches that strongly activated neurons in an intermediate model layer.^79^ The power spectrum of preferred patches for each blur level was compared with that of randomly sampled patches from untransformed ImageNet images (Figure S2). Full details are provided in the Supplemental Methods.

### Representational similarity analysis

Representational similarity analysis (RSA)^80^ was used to quantify correspondence between visual representations in the ventral visual stream (VVS) and DNNs. For both brain and model responses, pairwise similarity across the 36 objects was calculated using Pearson correlation, yielding a representational similarity matrix (RSM) that defined the representational geometry. Model–brain correspondence was then quantified as the Spearman correlation between brain and model RSMs.

#### Regions of interest

Regions of interest (ROIs) were defined using cytoarchitectonic parcellations from the Jülich atlas, normalized to age-appropriate templates for each age group (Figure S3). ROIs comprised V1 (hOc1), V2dv (hOc2), V3v (hOc3v), hV4 (hOc4v), VO1 (FG1),^81^ and FG4, a higher-level region in the fusiform gyrus which spans part of object-selective cortex in adults.^82^ For brevity, V2dv, V3v, and hV4 are referred to as V2, V3, and V4 throughout. For all ROIs, voxels from the left and right hemispheres were combined for analysis, as homotopic regions show similar patterns of visually evoked functional responses in young infants.^40^ Split-half noise ceilings for each bilateral ROI and age group are reported in Table S1.

#### Group-level representational geometry

Shared (group-level) visual representations were characterized for each age group in each ROI. Infant fMRI data were preprocessed using a custom pipeline described in detail in related work^41^ (see also Supplemental Methods; code is openly available at https://github.com/ClionaOD/foundcog_analysis). The pipeline included motion correction, normalization to age-appropriate templates and motion censoring. Volumes with a framewise displacement (FD) greater than 1.5 mm were excluded from analysis, as were runs in which more than half of volumes exceeded this threshold. Runs were also excluded if they were manually deemed of poor quality during the scanning session. A final sample of 101 2-month-olds (169 runs), 44 9-month-olds (64 runs), and 17 adults (51 runs) contributed to the group-level visual object representations. For each ROI, voxel-wise parameter estimates were obtained separately for each of the 36 objects. Pairwise Pearson correlations between multivariate object-response patterns were calculated across unique subject-run pairs, excluding comparisons between runs from the same participant, to create RSMs. The pairwise RSMs were then averaged and symmetry enforced to yield a group-level RSM, which characterized the shared representational geometry for each age group in each ROI.

#### Characterizing visual representations from DNN activations

To compute representational geometry for the DNNs, the FOUNDCOG objects at their maximum loom size (640 × 360 px) were placed on a neutral gray background and padded vertically with the same gray to center the object within a square image (640 × 640 px). Images were then resized to 224 × 224 px for presentation to the models. Activations were extracted following each convolutional ReLU layer of ResNet-50 (49 layers), and layer-wise representational geometry was calculated using Pearson correlations across responses to the 36 objects. This was repeated for test images across the same range of Gaussian blur used during training (σ = 0, 0.5, 1, 1.5, 2, 3, 4, or 6 px).

### Characterizing object-feature tuning across spatial scales

#### Spatial-scale tuning maps

A bootstrap resampling procedure was used to estimate model–brain correspondence for each age group and ROI. Group-level representational geometry was calculated across 1000 bootstrap resamples of subject-run pairs. Two independently trained models were available for each train-blur condition, and bootstrap resamples were balanced across model instances. For each age group and ROI, the median peak-layer correlation across bootstraps was calculated for every train–test blur combination, yielding a two-dimensional spatial-scale tuning map.

#### Developmental change in model–brain correspondence

To visualize developmental change, difference maps were constructed by subtracting the tuning map of the younger age group from that of the older age group. For each train–test blur combination, the bootstrap distribution of the developmental difference was calculated from paired peak-layer correlation estimates. Differences were considered reliable when the 95% bootstrap confidence interval excluded zero.

#### Estimating feature complexity: Peak-corresponding DNN layers across spatial-scale tuning maps

For each bootstrap and train–test blur combination, model–brain correspondence was calculated across DNN layers, and the layer yielding the highest correspondence was identified as the peak- corresponding layer. The corresponding model–brain correlation was used as the value for that cell of the spatial-scale tuning map. To summarize the peak-corresponding layer across the full tuning map, a weighted-average layer was calculated for each bootstrap by weighting the layer number associated with each cell by its corresponding model–brain correlation. The median and 95% bootstrap confidence interval of this distribution were used to summarize the peak-corresponding DNN layer for each age group and ROI (Figure 2F). Full calculation details are provided in the Supplemental Methods.

#### Quantifying tuning maturity: Similarity of infant and adult tuning patterns

To quantify the similarity of infant and adult tuning within each ROI, the covariance between the infant and adult tuning maps was calculated for each bootstrap and normalized by the variance of the corresponding adult tuning map. This provided a scaled measure of the extent to which infant tuning resembled the adult pattern, reflecting differences in tuning amplitude but insensitive to uniform offsets in model–brain correspondence, with a value of 1 corresponding to the adult reference value.

#### Testing for motion-related effects

To test whether residual motion contributed to developmental differences, the primary analyses were repeated using a more stringent run-level motion criterion. Runs with median framewise displacement (FD) > 0.5 mm were excluded, while timepoints in retained runs were censored as in the primary analysis when FD exceeded 1.5 mm. Sample characteristics for the resulting low-motion subset are reported in the Supplemental Methods.

### Correspondence with retinal-wave-trained models

Retinal-wave-trained models were obtained from the ReWaRD project.^50,54^ We compared ResNet-50 models trained on simulated retinal waves, trained on ImageNet-1K objects, pretrained on retinal waves and subsequently fine-tuned on ImageNet-1K, and an untrained randomly initialized network. FOUNDCOG object representations were extracted from all convolutional ReLU layers and compared with 1000 bootstrap samples of the brain data using the same peak-layer model–brain correspondence procedure described above. Model–brain correspondence was evaluated across the full range of test- blur levels used in the blur-trained model analysis and repeated using grayscale FOUNDCOG stimuli to test whether color information accounted for differences between models. Full model identifiers and implementation details are provided in the Supplemental Methods.

### Band-specific deep neural network models

#### Bandpass filtering

ImageNet images were decomposed into three spatial-frequency bands (low, mid, and high) using Butterworth filtering. The frequency boundaries were chosen with reference to the Gaussian blur levels used in the blur-trained models. The low band approximated frequencies retained following Gaussian blurring with σ = 4, while the high band approximated frequencies attenuated by Gaussian blurring with σ = 1. The mid band captured the intermediate range. The FOUNDCOG images used to evaluate model–brain correspondence were also decomposed into the same three bands.

#### Band-specific trajectories of model–brain convergence with visual cortex

As model–brain correspondence can emerge early during DNN training,^56^ we first characterized band-specific correspondence across an extended 60-epoch trajectory. Based on this preliminary analysis, we defined the first three epochs as the critical training window for the primary analysis (Figure S16; Supplemental Methods). Ten independently initialized instances of the network were trained for each spatial-frequency band. Across the training window, model representations were characterized approximately logarithmically as a function of batch number, providing denser sampling during the earliest stages of training while still spanning the full critical training window. Representations were additionally characterized for the untrained model and following the completion of each epoch.

At each sampled timepoint in the training trajectory (checkpoint), model representations for each spatial-frequency band were compared with visual cortical responses in V1–V4 for each age group using RSA. Each of the 10 independently initialized model instances was compared with the same 1000 bootstrap samples of the brain data. For each layer and bootstrap, model–brain correlations were averaged across model instances, after which the layer yielding the highest correspondence was selected as the peak layer. This yielded 1000 bootstrap trajectories of peak-layer model–brain correspondence for each age by ROI by spatial-frequency-band combination.

Band-specific differences in model–brain correspondence were directly evaluated at the endpoints of the critical training window: the untrained starting point and the final checkpoint of the three-epoch window. For each age and region, peak-layer model–brain correlations were compared pairwise across spatial-frequency bands using within-bootstrap differences, with correlations paired by bootstrap index. Band differences were considered reliable when the 95% bootstrap percentile interval of the paired difference excluded zero.

#### Characterizing learning dynamics with a logistic fit

For each bootstrap, a four-parameter logistic curve was fitted to the full trajectory of peak-layer model–brain correspondence, including the untrained model and all subsequent training checkpoints. Training time was transformed as log_10_(batches + 1), allowing the untrained model (0 batches) to be included in the fit. Fits were constrained to be non-decreasing (*k* ≥ 0). Additional constraints and usability criteria are described in the Supplemental Methods.

Letting *x* denote log-transformed training time and *y* denote model–brain correspondence, the trajectory was modeled as:

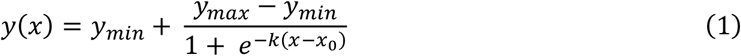

where *y_min_* is the lower asymptote, *y_max_* is the upper asymptote, *k* is the steepness of the curve and *x*_0_ is its inflection point (the midpoint of the curve).

Learning dynamics were subsequently summarized by three interpretable metrics: the ***midpoint of learning*** (the inflection point; *x*_0_) reflecting when the transition in model–brain correspondence occurred; the ***magnitude of learning*** (the fitted change in correspondence between the asymptotes; Δ*y* = *y_max_* − *y_min_*), and the ***trajectory steepness*** (the logistic steepness parameter; *k*), reflecting how abruptly model–brain correspondence changed around the inflection point.

##### Evaluating logistic fit

The quality of the logistic fit was evaluated, and a given age by ROI by band combination was considered sufficiently reliable for interpretation of the logistic parameters when at least 80% of the 1000 bootstrap fits met a set of usability criteria (Supplemental Methods).

##### Summarizing learning dynamics across bands

For age by ROI by band combinations that passed the logistic reliability criterion, parameter distributions were restricted to bootstrap fits that individually passed the usability criteria. For each combination, the median and 95% bootstrap percentile interval were calculated for the midpoint of learning, magnitude of learning, and trajectory steepness.

Within each age and ROI, spatial-frequency bands were compared using paired bootstrap differences for low versus mid, low versus high, and mid versus high bands. A contrast was considered reliable when the 95% bootstrap percentile interval of the paired difference excluded zero.

To assess the overall effect of training band across ages and regions, each retained age by ROI by band combination was first reduced to its median bootstrap parameter estimate. As the logistic reliability criteria resulted in some age by ROI combinations lacking an estimate for one spatial-frequency band, bands were compared using a Skillings–Mack test, a non-parametric extension of the Friedman test that accommodates incomplete blocks. Follow-up paired Wilcoxon signed-rank tests compared low versus mid, low versus high, and mid versus high using age by ROI combinations with estimates available for both bands. Pairwise p-values were corrected for multiple comparisons using the Holm procedure.

#### Band-dependent differences between infants and adults

To quantify developmental differences in relative band-specific model–brain correspondence at the end of the critical training window, peak-layer model–brain correspondence was compared between adults and infants within each ROI. Correlation coefficients were Fisher z-transformed before computing contrasts. For each age, ROI and bootstrap sample, z-values were demeaned across the three spatial-frequency bands, removing band-invariant differences in overall model–brain correspondence and isolating the relative pattern of correspondence across bands.

Adult–infant contrasts were then computed separately for each band by subtracting the relative band value for 2-month-olds or 9-month-olds from the corresponding adult value, with bootstrap samples paired by bootstrap index. This yielded a bootstrap distribution of the adult minus infant difference in relative band preference for each ROI and spatial-frequency band. The median and 95% bootstrap percentile interval were calculated for each contrast, and developmental differences were considered reliable when the 95% bootstrap interval excluded zero.

## Notes

https://github.com/dineenai/foundcog_coarse_fine

### Competing Interest Statement

The authors have declared no competing interest.

