## Supplementary material for "The development of coarse and fine feature tuning in infant visual cortex": Document S1. Supplemental information

### Supplemental Methods

#### Functional MRI setup and acquisition

Participants were scanned on a Siemens 3T MAGNETOM Prisma at the Trinity College Institute of Neuroscience, Trinity College Dublin. The infant scanning setup was designed to maximize comfort and minimize motion, following approaches developed for awake infant fMRI.[1,2] Detailed scanning logistics, comfort measures, and the visual presentation setup are described elsewhere.[3] Infants were positioned on a bead-filled vacuum pillow and wore Optoacoustics OptoActive II headphones, secured using inflatable cushions, for hearing protection and auditory stimulus delivery. Visual stimuli were projected directly onto the scanner bore above the infant's eyes using an Optoma UHD30 4K Ultra HD projector located in the technical room behind the scanner. Images were projected through an angled waveguide onto a mirror behind the head coil and then onto the bore. Stimuli were geometrically pre-distorted to correct for distortions introduced by the projection angle and bore curvature. Pre-distortion compressed each stimulus within the 640 × 360 px frame, so the maximum presented spatial frequency was estimated for each object from the bounding box of its non-transparent pixels in the pre-distorted image. The extent of the bounding box along the dimension that matched the display at maximum size (identified as the dimension with the greater pixel density, in pixels per degree) was divided by the visual angle of the corresponding display dimension (approximately 56° horizontally or 34° vertically), and the Nyquist limit was taken as half of the resulting pixels per degree. A member of the research team remained beside the infant throughout scanning, while infant state was monitored from the control room using an in-bore camera. The videos task was presented first, followed by the pictures task used in the present analyses. If the infant remained engaged, tasks were repeated. Each paradigm lasted approximately 5 minutes. Infant fMRI was acquired using the posterior array of a Siemens 64-channel head and neck coil. At 2 months, a 4-channel flex coil was additionally positioned above the infant's forehead to improve signal coverage of the frontal lobes. Functional images were acquired using a T2*-weighted multiband EPI sequence with a multiband factor of 4, TR = 610 ms, TE = 32 ms, flip angle = 40°, anterior-to-posterior phase encoding, 3 mm isotropic voxels, FOV = 192 × 192 mm, and a matrix of 64 × 64 × 36 voxels per volume. Adults were scanned using the full 64-channel head coil with a comparable multiband EPI sequence: multiband factor = 4, TR = 656 ms, TE = 30 ms, flip angle = 50°, left-to-right phase encoding, 3 mm isotropic voxels, FOV = 224 × 245 mm, and 40 slices per volume.

#### Functional MRI analysis and preprocessing

Infant fMRI data were preprocessed using a custom Nipype pipeline[4] developed by the FOUNDCOG team specifically for awake infant fMRI. The pipeline, including motion and data-quality procedures for the pictures task, is described in detail in related work using the same dataset[5] and is openly available at https://github.com/ClionaOD/foundcog_analysis. Briefly, for each functional run, EPI volumes were motion corrected using FSL MCFLIRT,[6] from which framewise displacement (FD) was estimated. Susceptibility-induced distortions were corrected using FSL TOPUP[7,8] with field maps acquired in opposite phase-encoding directions (AP/PA). Functional runs were then co-registered within participant using FSL FLIRT[6,9] with an affine transformation. Registrations were performed with 6, 9, or 12 degrees of freedom, with the best-fitting transformation selected for each participant following manual inspection, typically 12 degrees of freedom. Each participant's data were then normalized to common space using the age-appropriate NIHPD infant template (2–5-month or 8–11-month)[10] by application of a further affine transformation. Adult data were preprocessed using fMRIPrep[11] and normalized to the MNI152 template.[12]

For each functional run, a volumetric whole-brain general linear model (GLM) was fitted using Nilearn version 0.9.2.[13] A design matrix was constructed by modelling each of the 36 objects as a separate regressor and convolving each with a canonical hemodynamic response function. Nuisance regressors included the six motion parameters, a linear trend regressor, and a cosine drift model with a high-pass filter of 0.01 Hz. A motion threshold of 1.5 mm FD was used, with volumes exceeding this threshold censored from analysis.[14] Runs were excluded if more than half of the volumes exceeded this threshold or if the researcher attending the scan deemed the run to be of insufficient quality based on observation.[5] Overall, 15% of 2-month-old runs and 8% of 9-month-old runs were excluded on the basis of motion. A further 14% of runs at both ages were excluded because they were deemed poor quality by an attending researcher during the scanning session.[5]

Following preprocessing and run exclusion, 101 2-month-olds (169 runs), 44 9-month-olds (64 runs), and 17 adults (51 runs) contributed to the group-level visual object representations. Among included timepoints, median FD was 0.308 mm (IQR = [0.163, 0.637]) at 2 months and 0.267 mm (IQR = [0.161, 0.486]) at 9 months. For each ROI, a run-wise parameter estimate (beta) was calculated for each voxel and each object, yielding multivariate response patterns for subsequent representational similarity analysis. Voxels with unstable parameter estimates were excluded from analysis.

#### Split-half noise ceiling calculation

To estimate the noise ceiling for each ROI, split-half reliability was obtained by dividing the data from each cohort to create two group-level representational similarity matrices and calculating their correlation. The Spearman-Brown prophecy formula was then applied to adjust for the reduction in sample size that occurs when splitting the data (see Table S1).

#### Validating correspondence of ResNet-50 with infant visual responses

To confirm that the ResNet-50 architecture was suitable for our analyses, layer-wise representations from a standard ResNet-50, trained and tested on sharp (unblurred) images, were compared with visual responses along the developing VVS (Figure S4, right). The trained model showed robust layer-wise correspondence with visual responses across the VVS at all ages, with peak correlations comparable to or higher than those we previously reported using a supervised AlexNet.[5] Correspondence for untrained models is shown for comparison (Figure S4, left). We therefore used ResNet-50 for all spatial-scale tuning analyses.

#### Blur-trained DNN training implementation details

ResNet-50,[15] a standard convolutional neural network architecture, was trained on the ImageNet training dataset[16] using supervised learning. Training was implemented in Python 3.7.7 using PyTorch

1.7.1 and CUDA 10.1. Networks were trained for 60 epochs with a batch size of 256 using a standard ImageNet hyperparameter configuration, with stochastic gradient descent with momentum of 0.9 and an initial learning rate of 0.1, which decayed by a factor of 10 every 30 epochs. A standard ImageNet data augmentation procedure was used. Images were randomly cropped to 224 × 224 pixels and randomly flipped horizontally. For the blur-trained models, Gaussian blur was applied following the random crop using the torchvision GaussianBlur transform. The default kernel size of 9 pixels was used for σ = 0.5, 1, and 1.5 px. For larger levels of blur, the kernel size was increased to accommodate the larger filter while preserving image smoothness, with kernel sizes of 13, 17, 21, and 31 pixels used for σ = 2, 3, 4, and 6 px, respectively. Images were then converted to tensors and normalized using the ImageNet mean [0.485, 0.456, 0.406] and standard deviation [0.229, 0.224, 0.225].

#### Characterizing the spatial frequency of visual features extracted by DNNs trained on blurred images

For models trained on each level of blur, we identified a set of image patches from the model training dataset that led to high levels of activation in individual neurons from a chosen layer. Patches were identified from sharp images, which contain all spatial frequencies, thus serving as a broadband probe to test whether blur-trained models preferentially responded to image patches containing lower spatial frequencies relative to sharp trained models.

Patches were identified using the two-step process of Görgün and colleagues (https://github.com/adagorgun/VITAL).[17] We characterized the spatial frequency preference of an intermediate model layer (layer 31 – the output of residual stage 3 block 2, with 1024 neurons), analyzing responses from a random subset of 64 neurons. For each of these neurons, we identified the 10000 images that led to the highest levels of activation. From each image, 16 patches of size 64 × 64 px were extracted. The 16 patches from distinct images that led to the highest levels of activation for a given neuron were chosen as the preferred image patches for that neuron, yielding 1024 patches (16 patches for 64 neurons). As two models were trained per blur level, this process was repeated for each, yielding a total sample of 2048 preferred patches for the chosen layer trained on a given level of blur.

For each level of blur, the power spectrum was calculated and averaged across patches (Figure S2A). This was compared to the ImageNet baseline, which was calculated by taking a random sample of 16 patches from 500 images, giving a total of 8000 patches. To compare the model-wise spectra to the ImageNet baseline, the ratio of each power spectrum relative to baseline was calculated for each blur level (Figure S2B).

#### DNN object classification accuracy across train–test blur combinations

Classification accuracy was calculated for each train–test blur combination as the top-1 accuracy of each model on ImageNet validation images presented with each level of test blur (σ = 0, 0.5, 1, 1.5, 2, 3, 4, or 6 px). Accuracy was averaged across the two independently trained models for each train-blur condition, yielding a single classification-accuracy value for each train–test blur combination (Figure S8A).

#### Peak-corresponding DNN layer across the spatial-scale tuning map

For each bootstrap and train–test blur combination, model–brain correspondence was calculated across all DNN layers and the layer yielding the highest correspondence was identified as the peak-corresponding layer. This produced a layer map corresponding to the spatial-scale tuning map. Because the peak-corresponding layer varied across train–test blur combinations, a single summary value was calculated for each bootstrap by weighting the layer number associated with each cell by its corresponding model–brain correlation.

Letting $C$ denote the tuning map containing the peak-layer correlation $c$ for each train blur $tr$ and test blur $te$ combination, and $L$ the corresponding layer map containing the associated peak layer number $l$, the weighted-average layer for bootstrap $b$ was calculated as:

$$\frac{\sum_{tr,te} c_{tr,te} l_{tr,te}}{\sum_{tr,te} c_{tr,te}} = wal_b \tag{S1}$$

This yielded one weighted-average peak-layer estimate per bootstrap and therefore a distribution of 1000 map-level estimates for each age group and ROI. The median and 95% bootstrap confidence interval of this distribution were used to summarize the peak-corresponding DNN layer shown in Figure 2F.

### Testing for motion-related effects

To test whether residual motion contributed to the observed developmental differences, the primary analyses were repeated using a subset of runs meeting a more stringent motion criterion. Runs with a median framewise displacement (FD) > 0.5 mm were excluded, while timepoints within retained runs were censored as in the primary analysis when FD exceeded 1.5 mm. The resulting low-motion group-level RSMs included 97 runs from 2-month-olds (57% of the 169 runs included in the primary analysis; median FD = 0.23 mm), 51 runs from 9-month-olds (80% of 64 runs; median FD = 0.23 mm), and all 51 adult runs (median FD = 0.14 mm).

### Correspondence with retinal-wave-trained models

Retinal-wave-trained models were obtained from the ReWaRD project[18] via the open-access Zenodo repository.[19] We used ResNet-50 models trained for 90 epochs on simulated retinal waves (rwave-4096_resnet50_epoch90.pth),[20] trained on ImageNet-1K from random initialization (ImageNet-scratch-v1_resnet50_epoch90.pth), or pretrained on retinal waves and subsequently fine-tuned on ImageNet-1K (pt_rwave-4096_ft_ImageNet-v1_resnet50_epoch90.pth). A randomly initialized ResNet-50 was included as an untrained comparison.

FOUNDCOG object images were presented to each model, and activations were extracted from the 49 ReLU layers following convolutional operations. Layer-wise representational geometry was calculated and compared with 1000 bootstrap samples of the brain data for each age group and ROI. For each bootstrap, the layer yielding the highest model–brain correspondence was selected as the peak-corresponding layer, and the corresponding correlation was retained as the peak-layer model–brain correlation. The analysis was repeated using luminance-only grayscale versions of the FOUNDCOG stimuli to test whether differences between models could be explained by color information.

### Bandpass-trained models

#### Implementing the bandpass filter

Images were decomposed into three spatial-frequency bands (low, mid, and high) using Butterworth filtering. Butterworth filters were chosen as they provide a gradual transition around the frequency cutoff, reducing ringing (Gibbs) image artefacts compared with the hard cutoff of a brick-wall filter. This choice prioritized reducing artefacts and perceptual quality over a strict separation between bands.

Band boundaries were defined by two cutoff frequencies in cycles per pixel (cpp; where 0.5 cpp is the Nyquist frequency): 0.055 and 0.15 cpp. These cutoffs were empirically chosen to approximately match the frequency attenuation produced by Gaussian blur with σ = 4 and σ = 1 pixels, respectively (for details of the estimation procedure see below). These values were chosen as, in the previous experiment, application of Gaussian blur with a value of σ less than 1 resulted in DNNs that achieved high classification accuracy but were a poor fit for the early visual cortex and V4. This was therefore chosen as the high pass cutoff, while σ of 4 was chosen to approximate the lower frequency conditions in the blur-trained models.

Filtering was implemented using the skimage.filters.butterworth function[21] with a squared Butterworth filter (order = 3) and 40 pixels of padding to reduce boundary artefacts. The low band was obtained by low-pass filtering at 0.055 cpp, the high band by high-pass filtering at 0.15 cpp, and the mid band by low-pass filtering at 0.15 cpp followed by high-pass filtering at 0.055 cpp. Images were loaded as 8-bit unsigned integers (0–255). As mid- and high-pass filtering yields values centered on zero, each filtered image was scaled by its maximum absolute value to constrain values to the range [−1, 1], and then mapped back to [0, 255]. This mapping places zero at 127.5 (the midpoint of the 8-bit range), which appears as mid-gray. The low-band images did not require this re-centering. Values were then rounded, cast to 8-bit unsigned integers, and saved. Centering on mid-gray means that areas with little filtered content appear as a gray background, while lighter or darker regions indicate departures from this baseline. Using a fixed mid-gray baseline also kept the background level consistent across stimuli, helping ensure that comparisons were driven by filtered image content rather than differences in overall brightness.

#### Estimating equivalence between Gaussian blur and Butterworth cutoffs

Butterworth cutoff frequencies were estimated to produce frequency attenuation comparable to Gaussian blur with σ = 4 pixels (approximate low-pass boundary) and σ = 1 pixel (approximate high-pass boundary).

##### *Impulse image*

A single-channel 224 × 224 image containing a single white central pixel (112, 112) on a black background was generated. This 'impulse' stimulus provides a simple way to characterize each filter's effect on spatial frequencies, as the filter's frequency attenuation can be read out directly from the amplitude spectrum of the filtered impulse image.

##### *Candidate filtering and spectral summary*

The impulse image was filtered using Gaussian blur (σ = 4 or σ = 1) and a sweep of Butterworth filters spanning candidate cutoff frequencies. For each filtered image, we computed the Fourier transform and examined the Fourier magnitude spectrum. For comparison across filters, we extracted a 1D profile by taking a central horizontal slice of the 2D Fourier magnitude spectrum (values along $[x, 112]$), which summarises attenuation across spatial frequencies along one axis. We denote this 1D slice as the Fourier magnitude spectrum $A(f)$, where $f$ is spatial frequency along the horizontal axis (in cycles per pixel, cpp).

##### *Estimating equivalent low-pass boundary (σ = 4)*

To select the Butterworth cutoff corresponding to σ = 4, we compared the $A(f)$ from the Gaussian-filtered impulse image to the $A(f)$ produced by low-pass Butterworth filters across candidate cutoffs. The closest visual match in attenuation profile was obtained at 0.055 cpp.

*Estimating equivalent high-pass boundary (σ = 1)*

To select the Butterworth cutoff corresponding to the high-pass complement of a Gaussian blur with σ = 1, we constructed high-pass filtered images by subtracting the corresponding low-pass filtered impulse image from the original impulse image (high-pass = impulse − low-pass) for each candidate filter. We then computed Fourier magnitude spectra and extracted the same central horizontal slice, yielding $A(f)$ as above.

As Gaussian and Butterworth filters attenuate frequencies at different rates, the spectra were normalized to a common scale prior to comparison. Specifically, each 1D spectrum was normalized to its peak value ($max_f\{A(f)\}$) and converted to decibels relative to this peak (Equation S2).

$$A_{dB}(f) = 20log_{10}\left(\frac{A(f)}{max_f\{A(f)\}}\right) \quad (S2)$$

The $\sigma = 1$ Gaussian and Butterworth spectra were then compared at a common attenuation criterion of $-10$dB (relative to the peak) and the Butterworth cutoff whose spectrum most closely matched the Gaussian at this threshold was selected. This yielded a cutoff of 0.15 cpp.

#### Converting band boundaries to cycles per degree

The approximate cutoffs in cycles per degree of visual angle (cpd) were estimated. For the FOUNDCOG test images, bandpass filtering was applied in the DNN input space (224 × 224 px). Using cutoff frequencies of 0.055 and 0.15 cycles per pixel, the corresponding spatial frequencies were 0.055 × 224 = 12.32 cycles per image (cpi) and 0.15 × 224 = 33.6 cpi. When displayed to participants, the stimuli subtended approximately 56° of visual angle in width. Dividing cycles per image by degrees yields cutoff frequencies of approximately 12.32/56 = 0.22 cpd and 33.6/56 = 0.60 cpd.

#### Band-specific deep neural network models

As for the blur-trained DNNs, ResNet-50[15] models were trained on the ImageNet-1K dataset using supervised learning. Untrained ResNet-50 DNNs were randomly initialized using the standard torchvision implementation (Kaiming/He normal initialization for convolutional layers). Models were then trained on visual diets consisting of images from a single spatial-frequency band (low, mid, or high).

As decomposing images into frequency bands is computationally intensive, the ImageNet training and validation images were prefiltered into the three bands prior to training using the Butterworth filters defined above. Models were trained using a standard hyperparameter configuration with a batch size of 256, stochastic gradient descent with momentum (0.9) and an initial learning rate of 0.1, decayed by a factor of 10 every 30 epochs, where applicable. During training, standard ImageNet data augmentation was applied and images were randomly resized and cropped to 224 × 224 px. ImageNet images vary in size, with a median width and height of approximately 500 px and 375 px respectively. Filtering was applied before the random-resized crop, so variation in crop scale introduced corresponding variation in the effective spatial-frequency content of the final network inputs. Crops that zoomed in on a small portion of the prefiltered image were upsampled, shifting content toward lower frequencies, whereas larger crops were downsampled, shifting content toward higher frequencies. The effective frequency bands at the network input therefore varied across training samples, leading to partial overlap between bands. The low-, mid-, and high-frequency labels should therefore be interpreted as relative training constraints rather than perfectly fixed frequency ranges at the network input. Prefiltering nevertheless provided a practical means of imposing the intended frequency constraints at ImageNet scale while retaining standard ImageNet augmentation, including the scale variation introduced by random-resized cropping. This contrasts with the blur-trained models, for which images were first resized and cropped and subsequently blurred in the data loader. Images were then randomly flipped horizontally, converted to tensors, and normalized using the standard ImageNet channel-wise mean [0.485, 0.456, 0.406] and standard deviation [0.229, 0.224, 0.225]. Although the per-band channel-wise mean and standard deviation differed from the standard ImageNet values, we used the standard ImageNet normalization values for all models to maintain consistent input scaling and facilitate direct comparability across training conditions.

#### Classification accuracy of DNNs trained on different frequency bands

For each band, a single DNN instance was trained to 60 epochs. At the end of each epoch, categorization ability was determined for each model by calculating its top-1 classification accuracy in response to the ImageNet validation images.

#### Logistic fit and reliability criteria

For each bootstrap, a logistic function was fit to the trajectory of peak-layer model–brain correspondence. A fit was considered usable for parameter interpretation if optimization converged successfully, all fitted parameters were finite, the estimated midpoint fell within the critical training window, and the fitted magnitude parameter ($\Delta y$) fell within the interval $0 \leq \Delta y \leq 2$, corresponding to the maximum possible difference between two correlation coefficients. Logistic trajectories were constrained to be non-decreasing ($k \geq 0$), and an upper numerical bound of $k = 200$ was imposed for optimization stability. Fits reaching this upper bound were recorded for diagnostic purposes but were not excluded.

An age by ROI by band combination was considered sufficiently reliable when at least 80% of the 1000 bootstrap fits were usable. Sensitivity to this criterion was assessed using thresholds of 70% and 90%. The inclusion pattern was unchanged at 70%, while the more stringent 90% threshold resulted in additional exclusions for the mid-spatial-frequency band in infants. Median adjusted coefficient of determination ($R^2_{\mathrm{adj}}$) was additionally reported as a measure of fit quality but was not used as an exclusion criterion. Logistic fits were also compared with a simpler log-linear alternative, with the logistic model providing a better fit in the majority of comparisons.

#### Defining the critical training window from preliminary trajectories of model–brain convergence in visual cortex

Each training epoch comprised approximately 1.28 million images, corresponding to 5005 batches at a batch size of 256 images. To estimate the approximate learning trajectory while retaining high temporal resolution during early training, a single model instance for each band was trained for 60 epochs. Model accuracy was evaluated at the end of each epoch to verify that each band contained learnable object information. Model representations were sampled throughout training on a logarithmic scale, such that sampling was densest early in training while still spanning the full 60-epoch trajectory.

At each sampled checkpoint, the bootstrap distribution of peak-layer model–brain correlations was estimated and a logistic function was fit separately to each bootstrap trajectory. For this preliminary analysis, the fitted midpoint of model–brain convergence was the primary parameter of interest, as its purpose was to identify the period over which the principal representational changes occurred. Using the reliability criteria defined above, 25 of the 36 age-by-band-by-ROI trajectories were sufficiently reliable for interpretation.

Across these reliable trajectories, the median fitted midpoint was 0.265 epochs, with age by ROI by band median midpoints ranging from 0.002 to 1.149 epochs. Moreover, the full 95% bootstrap interval for the midpoint fell within the first three epochs for 23 of the 25 reliable trajectories. We therefore defined the first three epochs of training as the critical training window for the primary analysis.

This preliminary analysis incorporated variation across participants through bootstrap resampling but used only a single trained model instance per spatial-frequency band. The primary analysis was therefore repeated across 10 independently trained model instances per band to additionally account for variability in model training, which is particularly relevant during the early stages of learning.

### Supplemental Figures

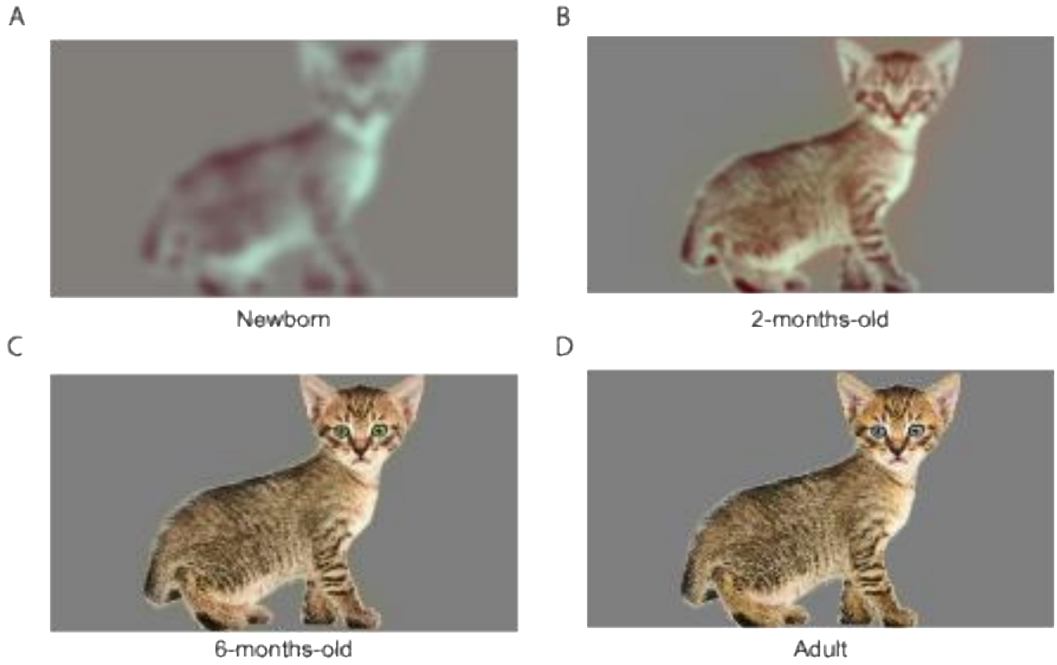


**Figure S1. Simulated appearance of a sample object on the FOUNDCOG display across the first postnatal year.**
Visual acuity and color vision develop rapidly over the first postnatal year.[22-27] Consequently, the test-time appearance of visual stimuli also changes. To estimate the appearance of our stimuli at 2 months corrected gestational age, we used the TinyEyes simulator (https://github.com/wadelab/VischeckTinyeyes), which simulates infant vision given the display parameters at test time. Given our display parameters (objects spanning up to 29 cm in width, presented approximately 27 cm from the infant's eyes, yielding a maximum spatial frequency of approximately 4.9 cpd), the objects were estimated to remain clearly identifiable at 2 months (B). The estimated appearance of the same stimulus to a newborn (A), a 6-month-old (C) and an adult (D) is shown for comparison.

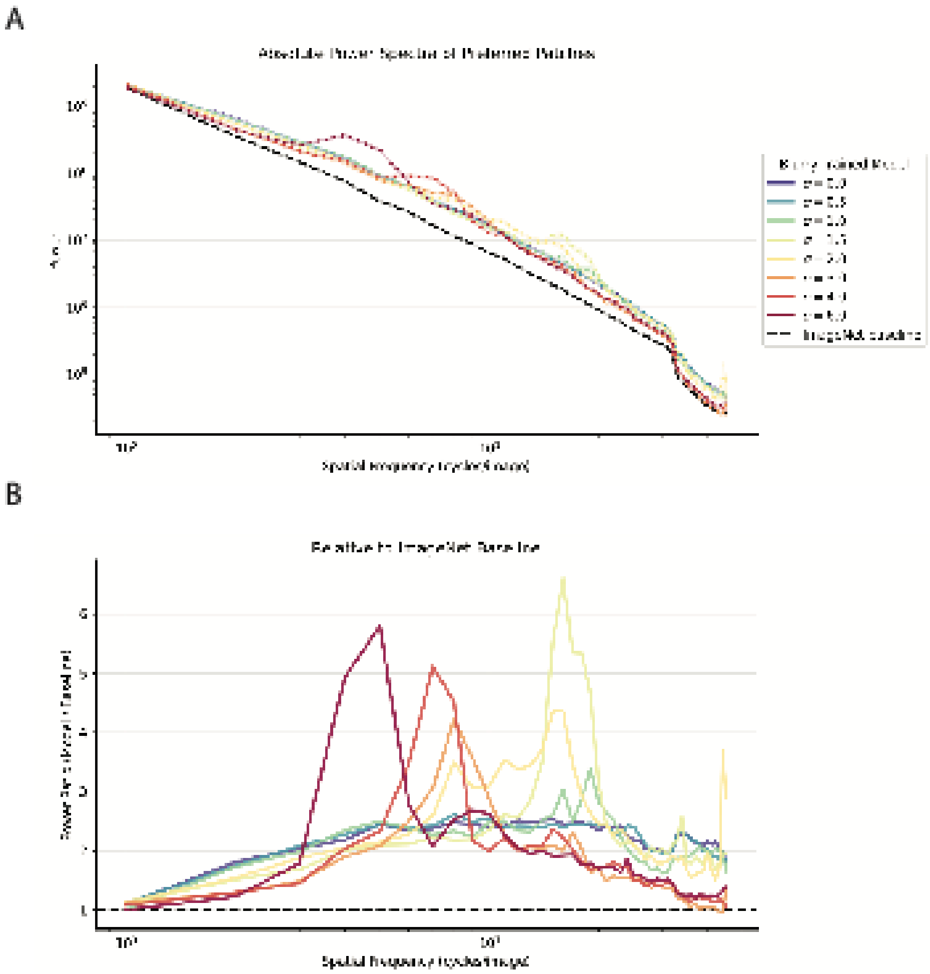

**Figure S2. DNNs trained on blurrier images preferred coarser object features.**

For ResNet-50 models trained on each level of blur (σ = 0–6), we identified a set of image patches (n = 2048) from the standard, sharp model training dataset (σ = 0) that led to high levels of activation in neurons in an intermediate model layer (layer 31). The average power spectrum of the preferred patches for each blur level was calculated and is depicted in (A), with the error band showing the 95% CI across 1000 bootstrap resamples. The baseline power spectrum was estimated from a random sample of patches (n = 8000) from the same dataset (dashed black line). As expected for natural images, the baseline power spectrum showed an approximately power-law decay with spatial frequency (typically found to be close to $1/f^2$ ), as evidenced by a near-linear trend on the log–log plot.[28,29] The ratio of the power spectrum for the blur-trained models relative to this baseline is depicted in (B).

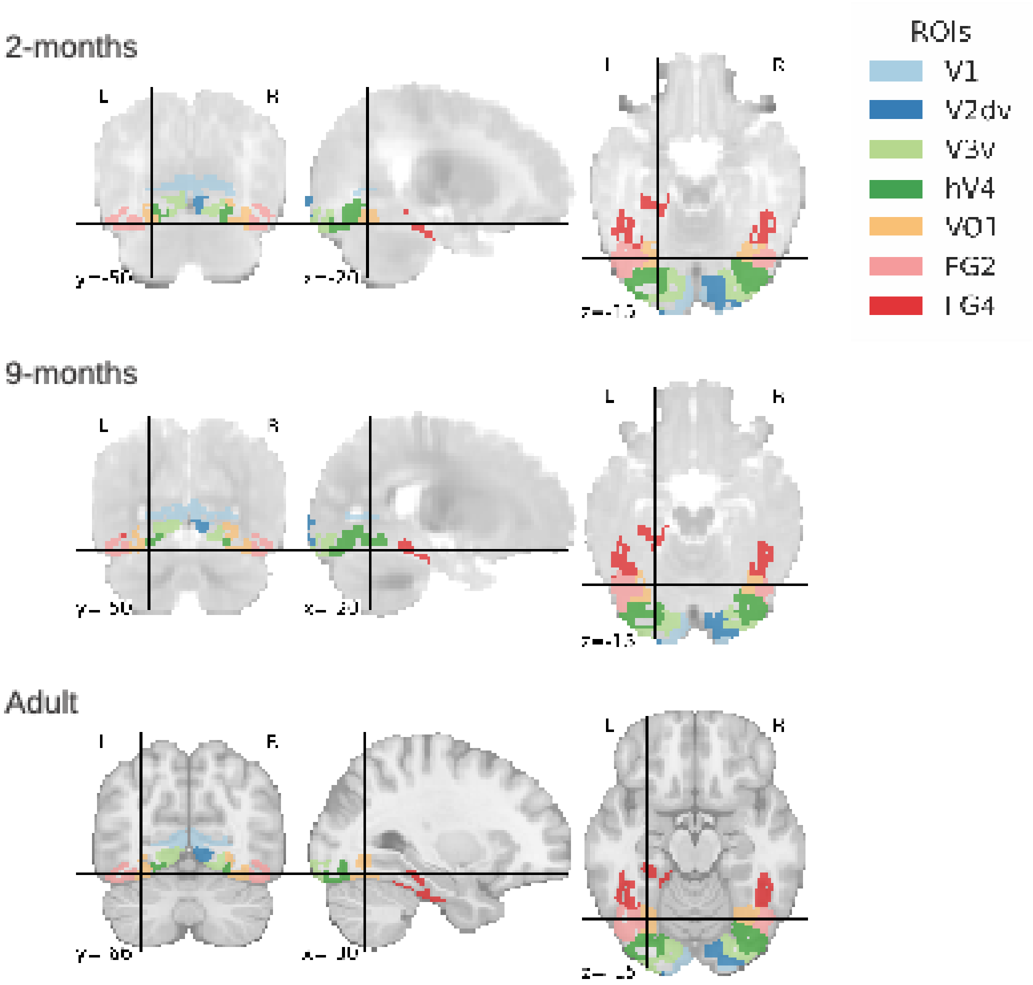


**Figure S3. Ventral visual stream regions of interest for which tuning to visual object features was characterized.**

Regions of interest (ROIs) were defined using cytoarchitectonic parcellations from the Jülich atlas,[30,31] which have been previously cross-validated as overlapping with functional regions of the ventral visual stream in adults ( see Methods). ROIs were normalized to age-appropriate templates and are visualized for each age group on the corresponding template. An NIHPD template was used for infants[10] and the MNI template was used for adults.[12]

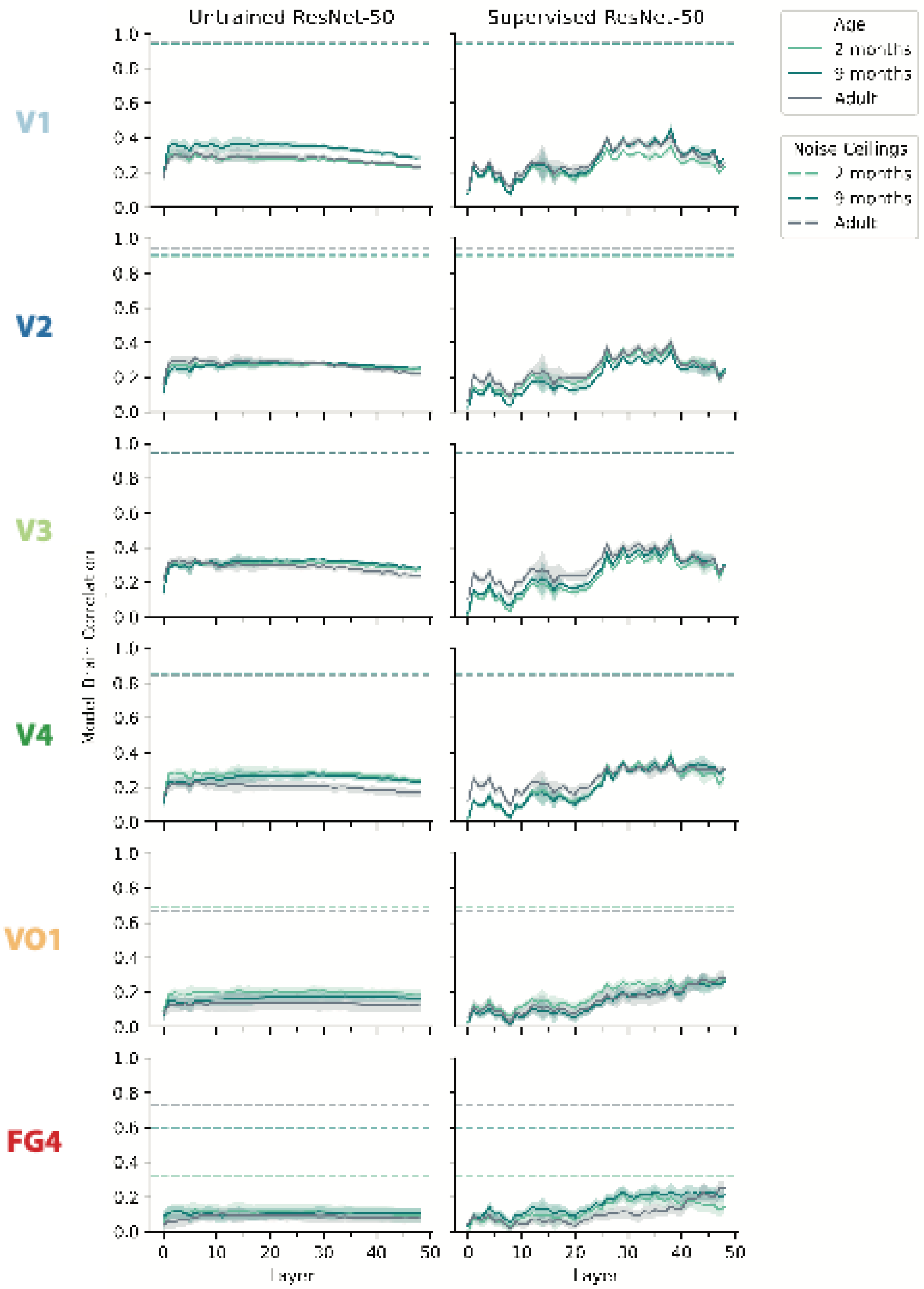


**Figure S4. ResNet-50 captures developing object responses along the ventral visual stream.** Layer-wise correspondence between developing visual responses along the ventral visual stream (VVS) and representations from ResNet-50. Panels on the left show model–brain correspondence for untrained models, whereas panels on the right show correspondence after supervised training on ImageNet. For each age group and region, line plots depict the median model–brain correlation across 1000 bootstrap resamples, with error bands showing the 95% confidence interval. Two model instances were used, corresponding to two random initializations; 500 bootstraps were compared to each instance, giving 1000 bootstraps per condition. Dashed lines indicate the age-specific split-half noise ceiling for each region.

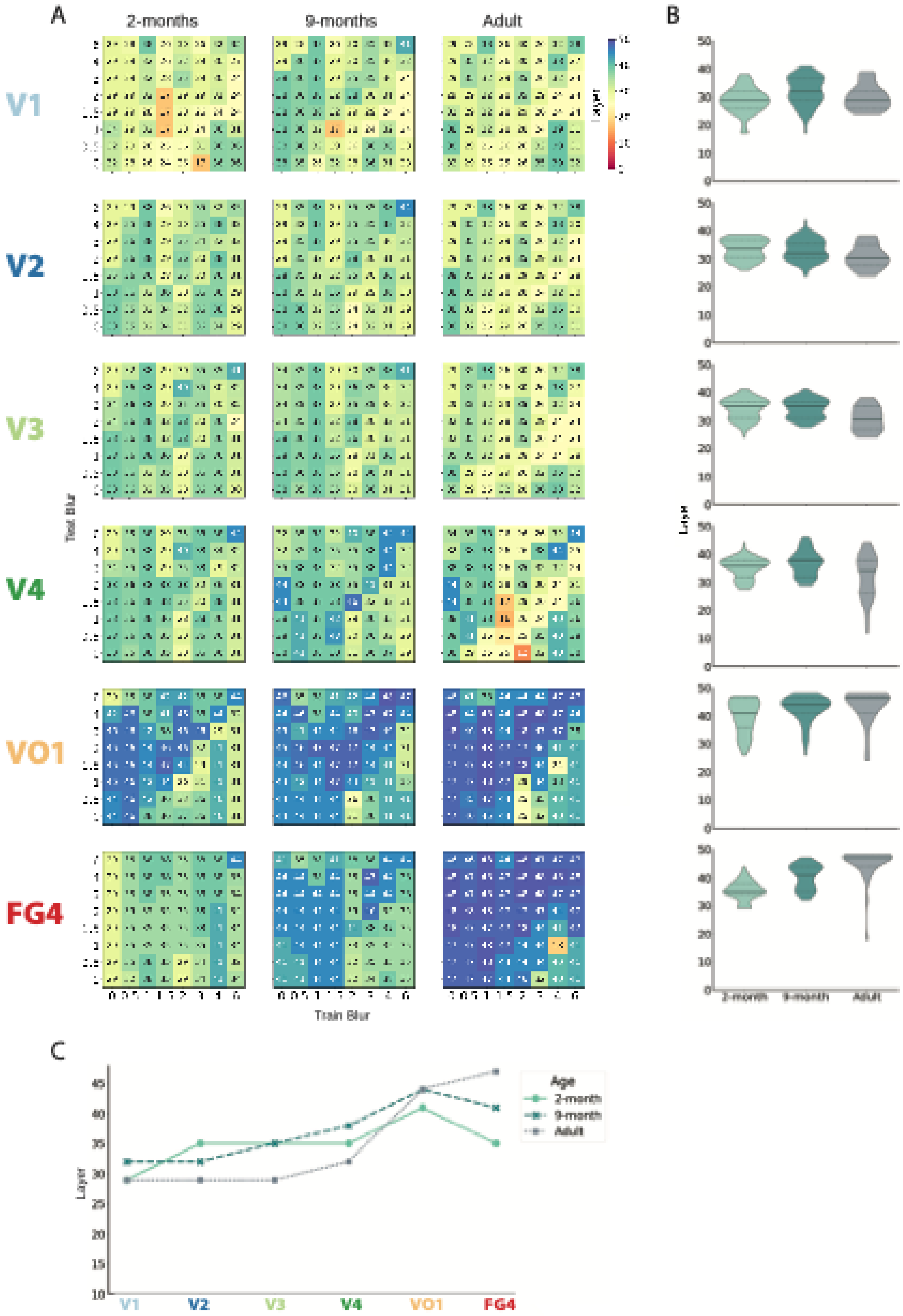


**Figure S5. Peak-corresponding DNN layers across spatial-scale tuning maps along the developing ventral visual stream.**

Early visual regions (V1–V4) were best captured by intermediate DNN layers, whereas VO1 was better captured by later layers across development. FG4 showed a developmental shift in its peak-corresponding layers, with 2-month-old responses better captured by earlier layers, consistent with representations of lower feature complexity at this age. For each ROI, age group, train–test blur combination, and bootstrap resample, model–brain correspondence was calculated across DNN layers and the layer yielding the highest correspondence was identified as the peak-corresponding layer. (A) Median peak-corresponding layer across 1000 bootstrap resamples for each train–test blur combination. (B) Distribution of the median peak-corresponding layers shown in (A) across train–test blur combinations for each ROI and age group. Thick black lines indicate the median and dashed lines the interquartile range. (C) Median peak-corresponding layer across train–test blur combinations, providing a descriptive summary of the distribution shown in (B).

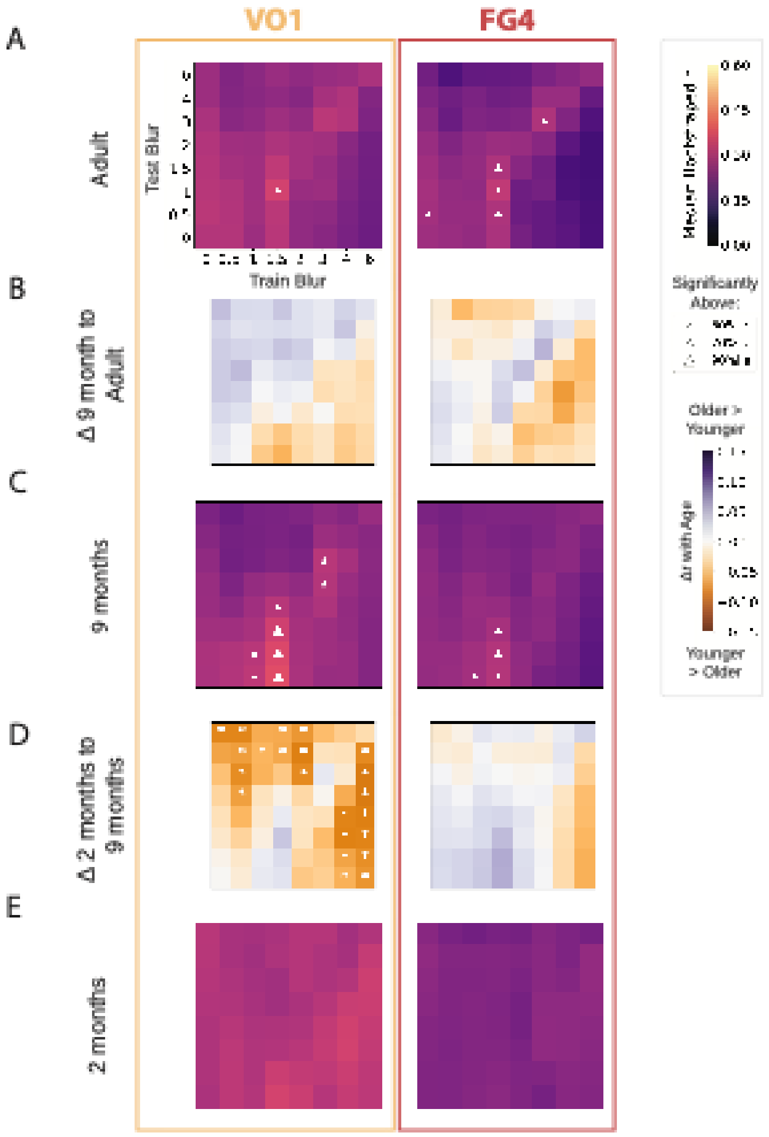


**Figure S6. Spatial-scale tuning maps for later visual regions.**
Spatial-scale tuning maps for later ventral stream regions, VO1, an intermediate visual region, and FG4, a region in category-selective cortex, are shown for adults (A), 9-month-olds (C), and 2-month-olds (E). Tuning maps were calculated as in Figure 2A, 2C, 2E. Developmental differences in tuning patterns between 2 months and 9 months (D), and 9 months and adults (B) were calculated by subtracting the tuning map of the younger age from that of the older age, as described for Figure 2B and 2D.

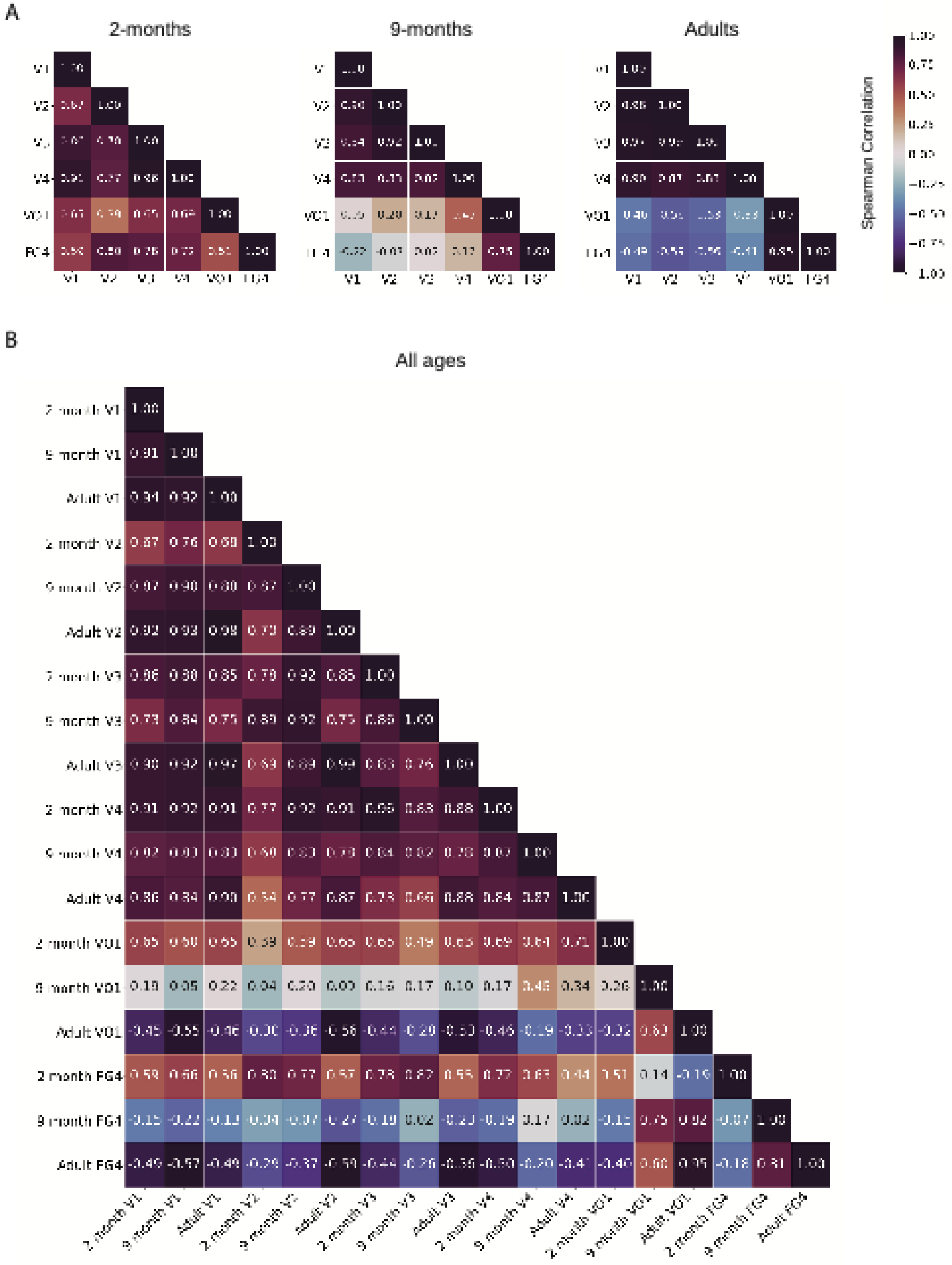


**Figure S7. Similarity of spatial-scale tuning maps along the ventral visual stream across development.**
(A) Spearman correlations between tuning maps within each age group. At 2 months, tuning in VO1 and FG4 was more similar to that in V1–V4 than at older ages. By 9 months, tuning in these later VVS regions had become more distinct from V1–V4, and by adulthood the tuning patterns in earlier and

later visual regions were negatively correlated. (B) Spearman correlations between tuning maps across age groups highlight the similarity of tuning patterns within V1–V4 across development.

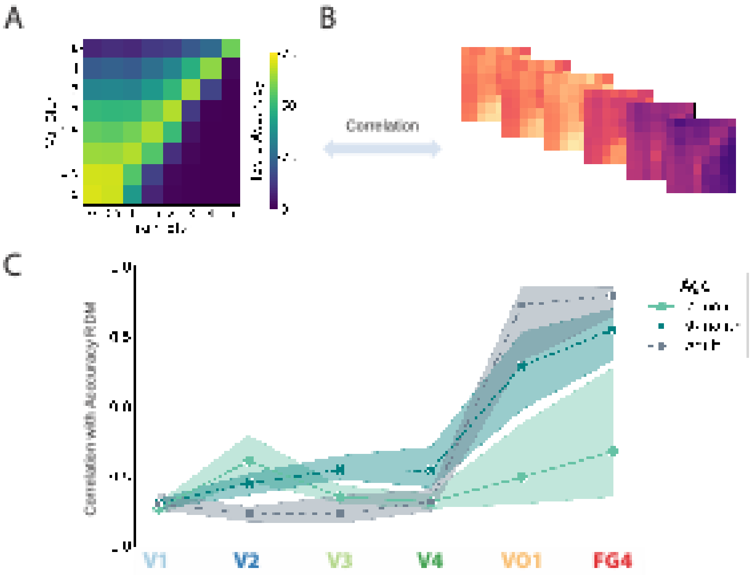


**Figure S8. Correlation between tuning maps and blur-dependent model classification accuracy.** Classification accuracy was calculated for each train–test combination of blur by calculating the top-1 classification accuracy of each model in response to the ImageNet validation images (val) and averaging across model instances (A). The matrix of blur-dependent model accuracy was correlated with tuning maps along the VVS (B). High levels of similarity would indicate that a region at a given age corresponded well with models with high classification accuracy. Median bootstrapped correlation with 95% confidence interval between tuning maps and the blur-dependent accuracy map along the VVS across development (C). Tuning maps in early visual cortex were negatively correlated with model accuracy at all ages, whereas later VVS regions became progressively more correlated with model accuracy across development.

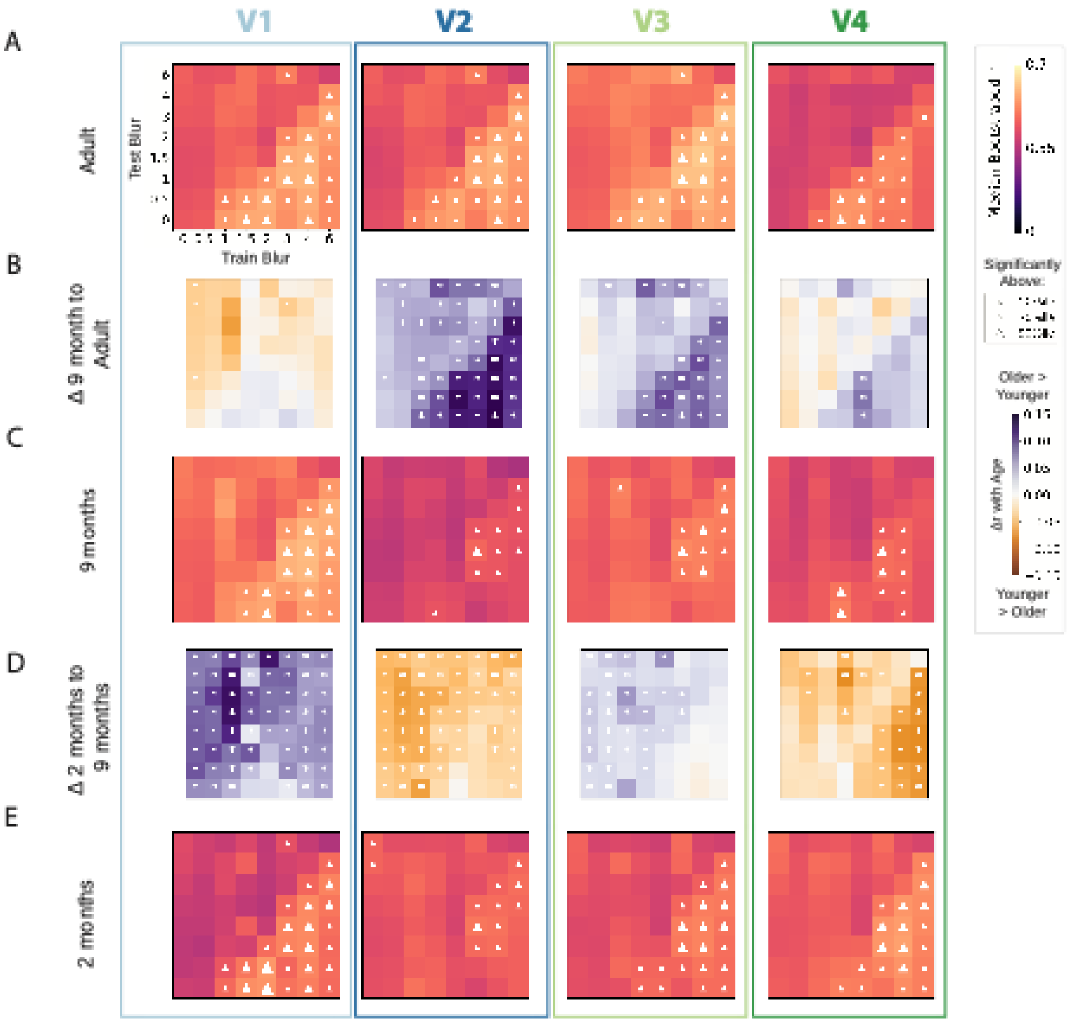


**Figure S9. Tuning to coarse and fine object features along the ventral visual stream scaled by the noise ceiling.**
Spatial-scale tuning maps for the early visual cortex of adults (A), 9-month-olds (C) and 2-month-olds (E) scaled by the appropriate noise ceiling (as defined in Table S1). Tuning maps were calculated and are depicted as described for Figure 2A, 2C, 2E. Developmental differences in the scaled tuning maps between 2 months and 9 months (D), and 9 months and adulthood (B), were calculated by subtracting the tuning map of the younger age group from that of the older age group, as described for Figure 2B and 2D.

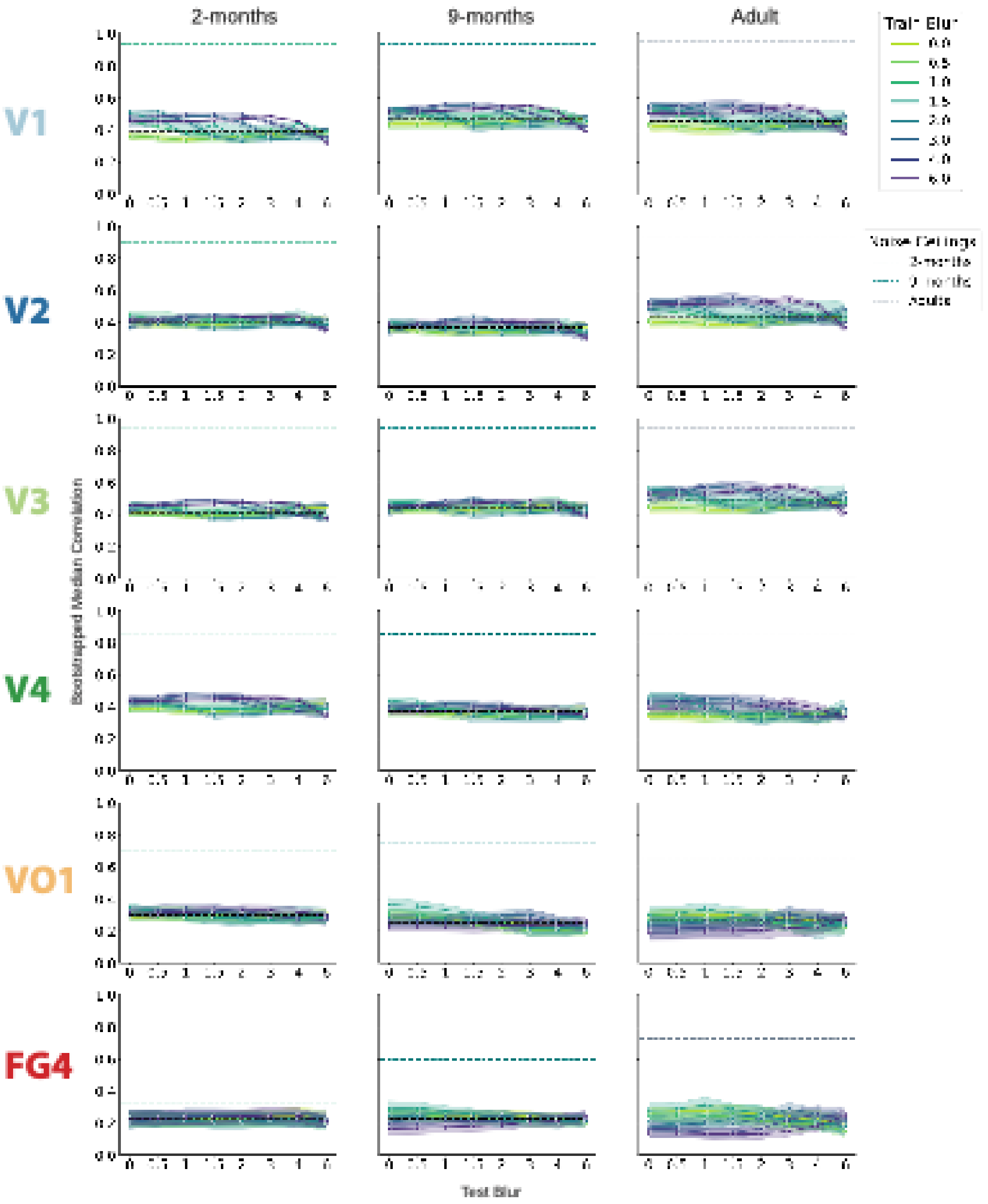


**Figure S10. Reliability of spatial-scale tuning across the developing ventral visual stream.** Alternative visualization of the model–brain correspondences characterized by the spatial-scale tuning maps in Figure 2 and Figure S6. Each colored line shows the bootstrapped median correlation value of the layer yielding the highest correlation with the brain (y-axis) for a given value of training blur (hue) across the range of test blurs (x-axis). The error band depicts the 95% confidence interval of the correlation across bootstrap resamples. The thick black dashed line depicts the median peak-layer correlation across all bootstraps for all combinations of train and test blur. The dashed green, teal and gray lines denote the split-half noise ceiling for each region for 2-month-olds, 9-month-olds and adults respectively, setting an upper bound on model correspondences. Correlation values were not scaled by the noise ceiling as this would inflate estimates in regions with low split-half reliability, such as FG4 at 2 months.

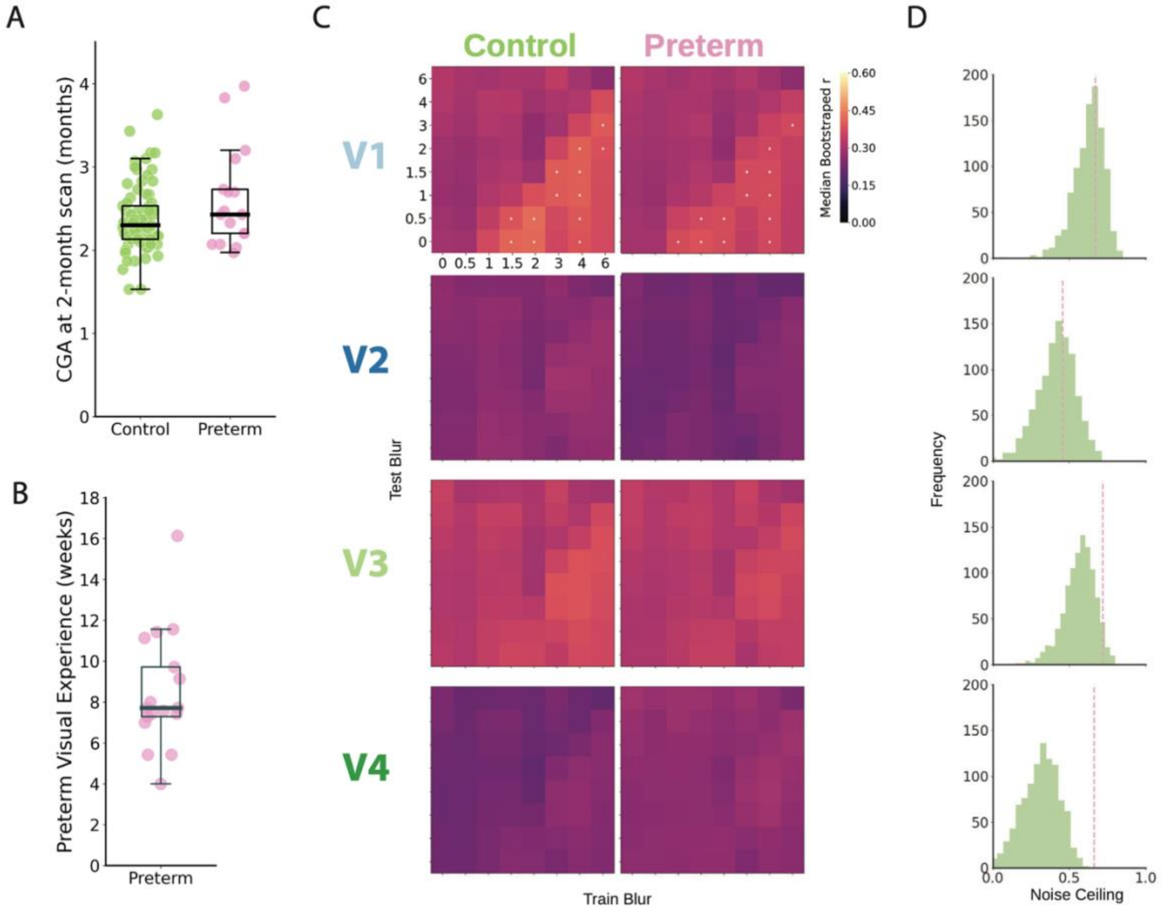


**Figure S11. Spatial-scale tuning at 2 months was similar in preterm-born and term-born infants.**
To investigate whether cumulative early visual experience was associated with spatial-scale tuning, a subgroup of preterm-born infants who had spent time in the neonatal intensive care unit (NICU; n = 17) was compared with bootstrap samples of run-matched term-born controls. The groups were similar in corrected gestational age (CGA) at scan (A), but the preterm-born infants had been born substantially earlier and had therefore experienced a median of 7.71 additional weeks of postnatal visual experience by the time of scanning (IQR = [7.29, 9.71] weeks; B). Spatial-scale tuning was highly similar between groups, with no qualitative differences in the tuning maps (C). The split-half noise ceiling for the preterm-born group (dashed pink line) fell within the distribution obtained from bootstrap samples of run-matched controls (green distribution) in V1 and V2. In V3 and V4, the split-half noise ceiling was higher in the preterm-born group, suggesting greater consistency of object-response patterns across infants in these regions (D).

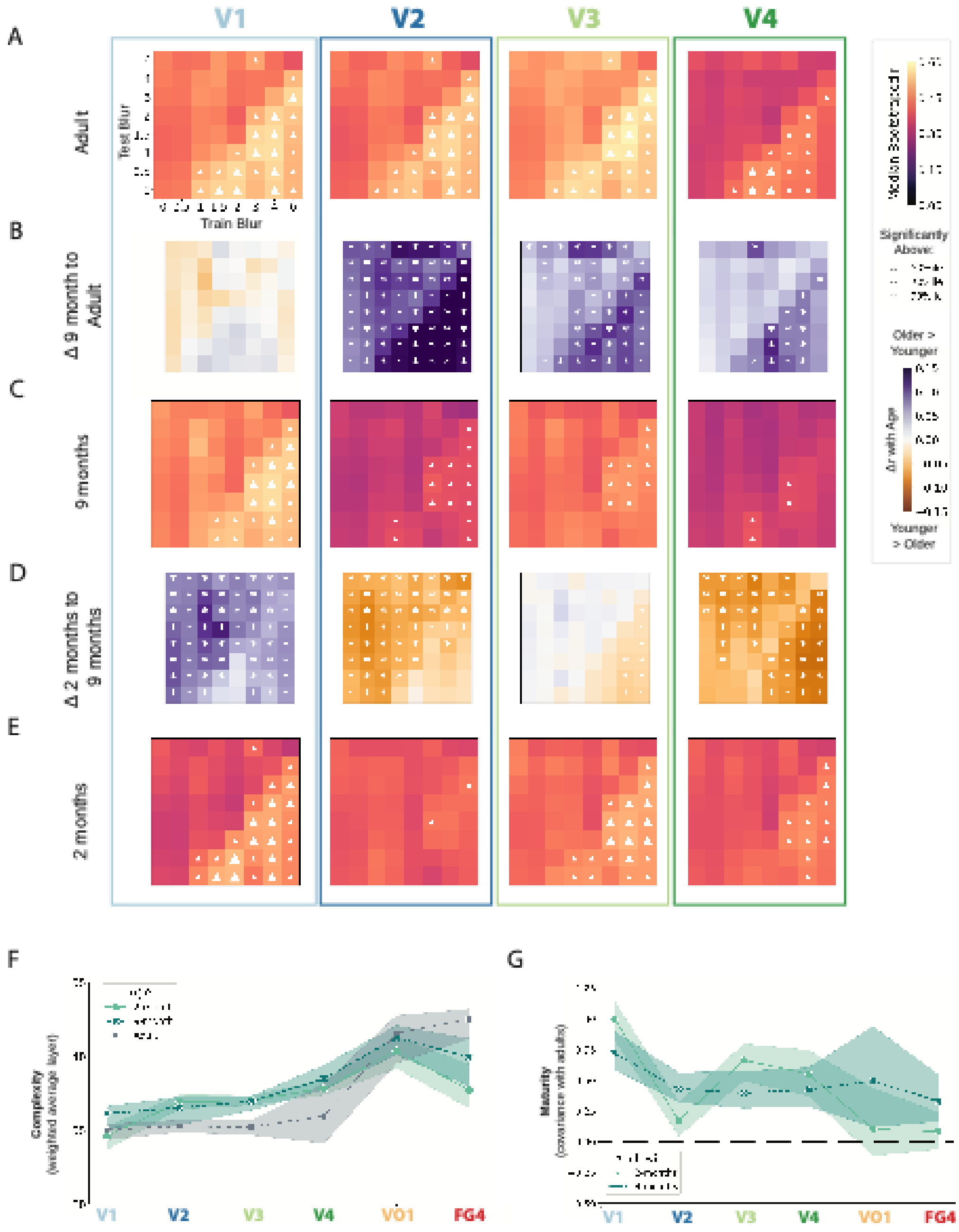


**Figure S12. Replication of spatial-scale feature tuning using a low-motion subset of infant runs.** The tuning patterns shown in Figure 2 were replicated when analyses were repeated using a low-motion subset of infant runs (see Methods). Tuning to object features across spatial scales is shown for adults (A), 9-month-olds (C), and 2-month-olds (E), with infant runs restricted to those with median FD below 0.5 mm. Difference maps (B, D), weighted average peak-layer estimates (F), and maturity analyses (G) were computed as in Figure 2. Overall, the developmental pattern was similar to that observed in the full dataset, with V1 appearing relatively mature by 2 months and later regions showing more protracted development. Analysis details and plotting conventions are consistent with those described for Figure 2.

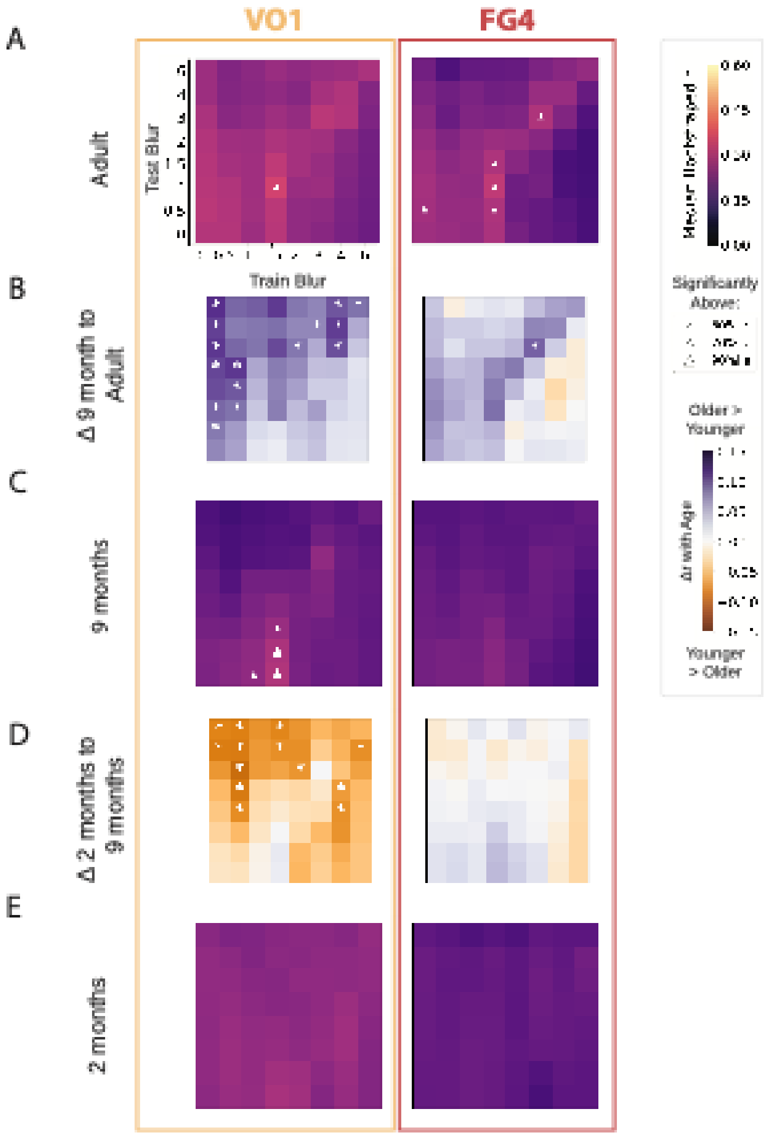


**Figure S13. Replication of spatial-scale tuning for later visual regions using a low-motion subset of infant runs.**
Repeating the analysis shown in Figure S6 with a low-motion subset of infant runs (see Supplemental Methods) produced a similar pattern of results.

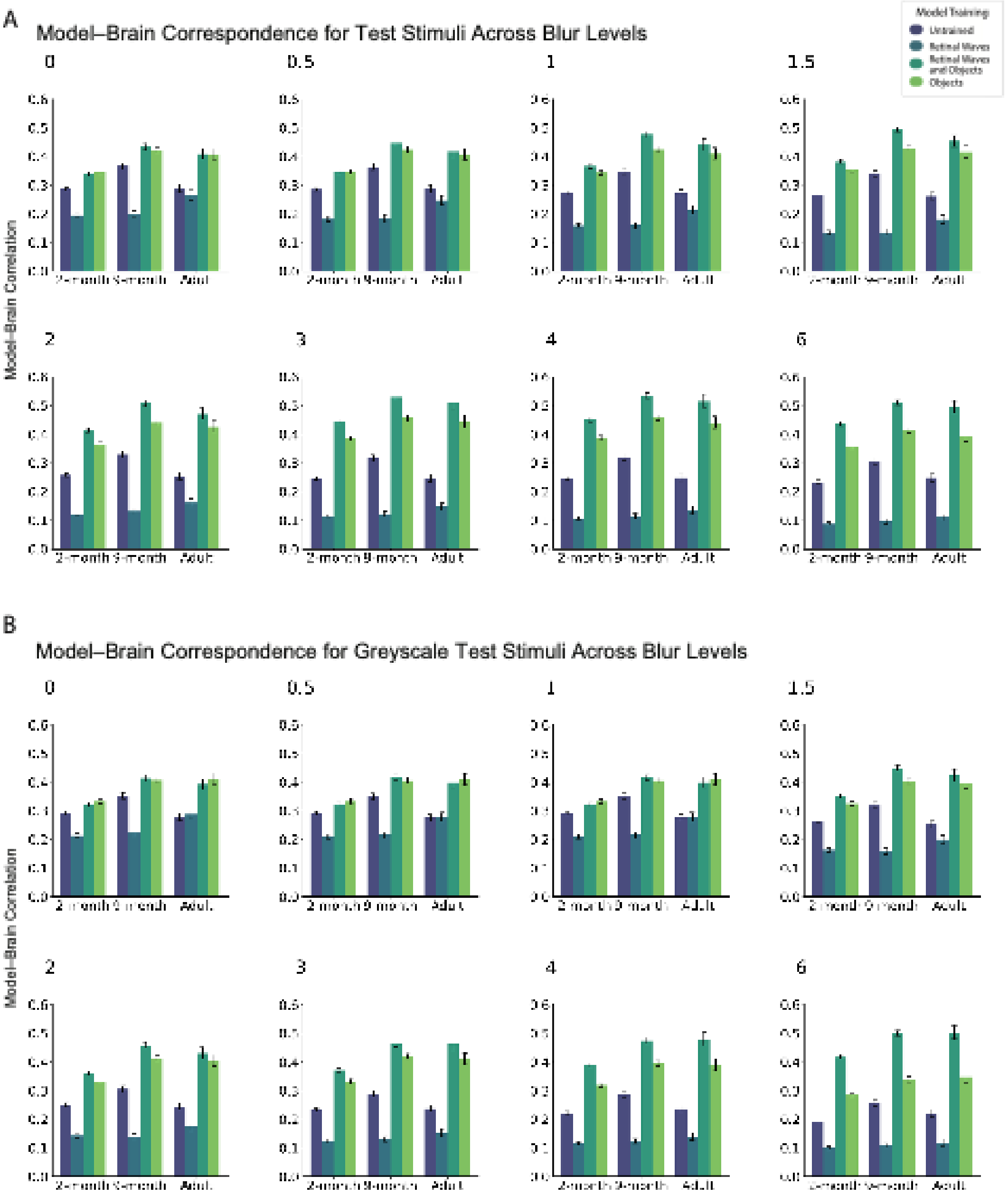


**Figure S14. Correspondence between V1 and retinal-wave-trained models, tested on images across a range of blurs.**

At all ages, the retinal-wave-trained model was a poorer model of V1 than the object-trained model. This suggests that, as early as 2 months, V1 representations are better captured by features learned through object training than by features learned from retinal waves. Each graph depicts the model-wise results when tested on the FOUNDCOG stimuli at the specified level of test blur. For sharp test images, pretraining on retinal waves conferred no advantage over standard object training. At higher levels of test blur, however, pretraining on retinal waves resulted in increased representational correspondence with V1 (A). To determine whether color might be driving the differences between retinal-wave-trained and object-trained models, the analysis was repeated with grayscale versions of the test stimuli. Grayscale images produced a similar pattern of results (B), suggesting that color is not driving the observed results.

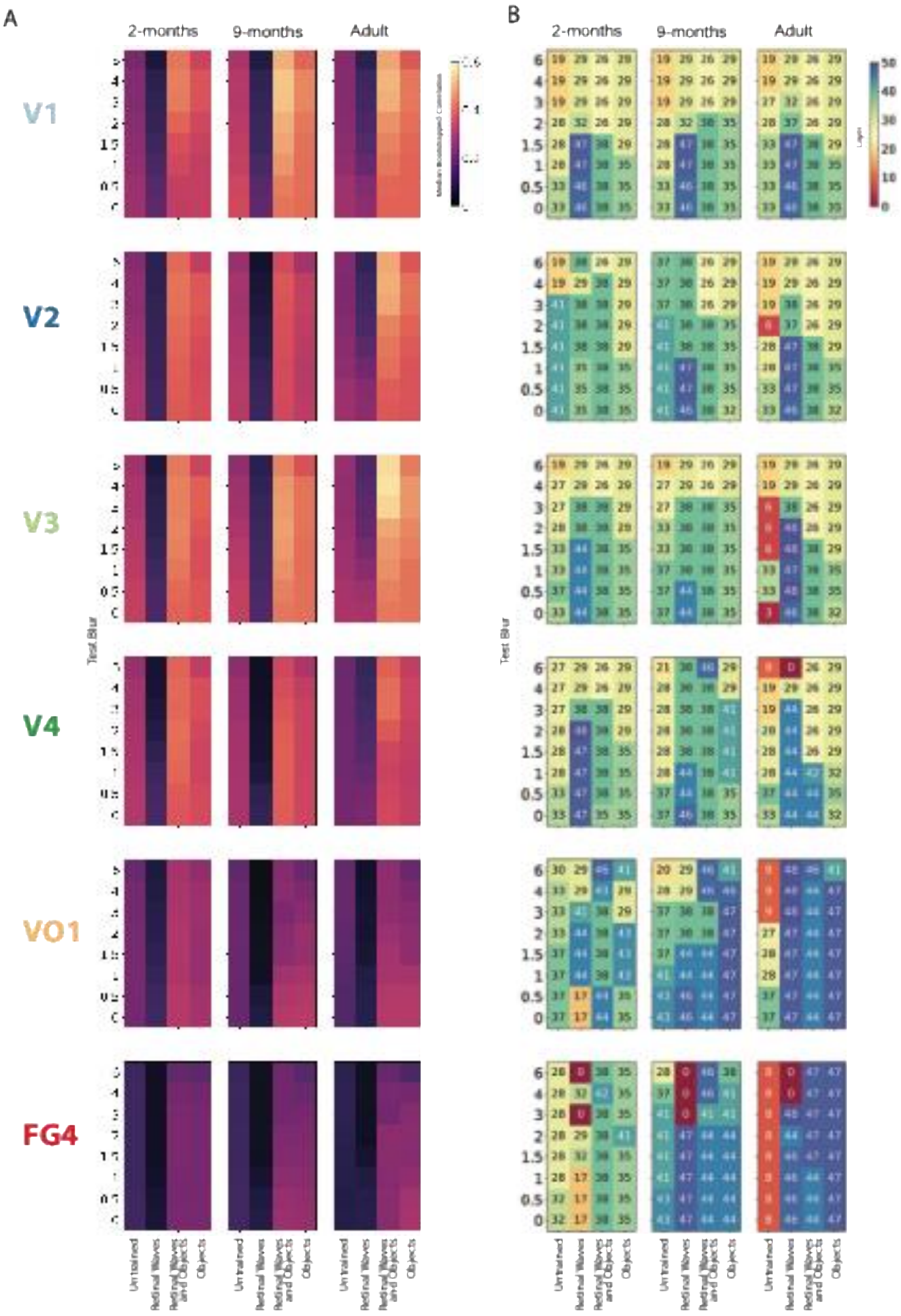


**Figure S15. Model–brain correspondence for retinal-wave-trained models along the ventral visual stream.**

Across early visual cortex and V4, the pattern of model–brain correspondence was similar to that observed in V1 (Figure S14A) for both models pretrained on retinal waves and subsequently trained on objects and models trained only on objects. In later ventral visual regions, VO1 and FG4, retinal-wave pretraining no longer increased model–brain correspondence relative to object training alone.

(A) For each model, test-blur level, age group, and ROI, layer-wise model–brain correspondence was calculated and the layer yielding the highest correspondence (peak layer) was identified. The bootstrap median correlation at the peak layer is shown. (B) The corresponding bootstrap median peak-layer number is shown for each model by test-blur combination, analogous to the peak-layer estimates shown in Figure S5A.

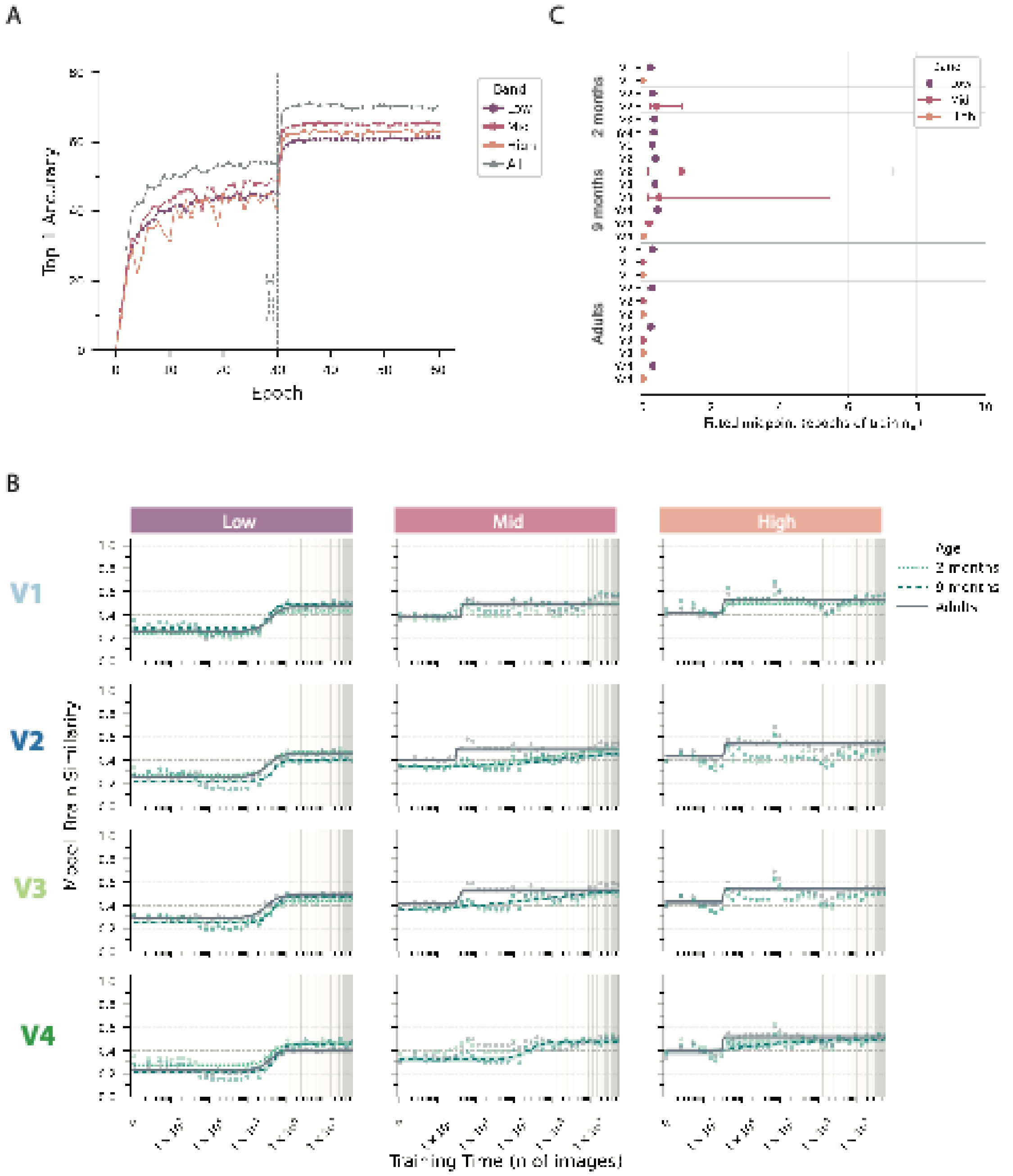


**Figure S16. Defining the critical training window for model–brain convergence.** Preliminary characterization of band-specific model–brain correspondence across an extended 60-epoch training trajectory. (A) One network instance per spatial-frequency condition was trained for 60 epochs on low-, mid-, high-, or all-frequency ImageNet images. Top-1 categorization accuracy was evaluated at the end of each epoch on the corresponding ImageNet validation set, confirming that each spatial-frequency band contained sufficient information to support object learning. (B) Model representations were sampled throughout training at approximately logarithmically spaced batch checkpoints, providing dense sampling early in training while still spanning the full 60-epoch trajectory. At each sampled checkpoint, model representations were compared with visual representations in V1–V4 in 2-month-olds, 9-month-olds, and adults. For each age by ROI by band combination, peak-layer model–brain correspondence was estimated across 1000 bootstrap resamples of the brain data and normalized by the age- and ROI-specific noise ceiling. Points show the bootstrap median at each sampled checkpoint. Logistic functions were fitted separately to each bootstrap training trajectory. An age by ROI by band trajectory was considered sufficiently reliable for interpretation when at least 80%

of its bootstrap fits met the specified usability criteria. For reliable trajectories, the median logistic fit and 95% bootstrap interval are shown. (C) The inflection point of each reliable logistic fit was used to estimate the midpoint of learning in the preliminary training trajectory. Across reliable trajectories, the median fitted midpoint was 0.265 epochs, with age × ROI × band medians ranging from 0.002 to 1.149 epochs. These results showed that the principal changes in model–brain correspondence were concentrated very early in training and motivated defining the first three epochs as the critical training window for the main analysis. The dashed gray line marks three epochs.

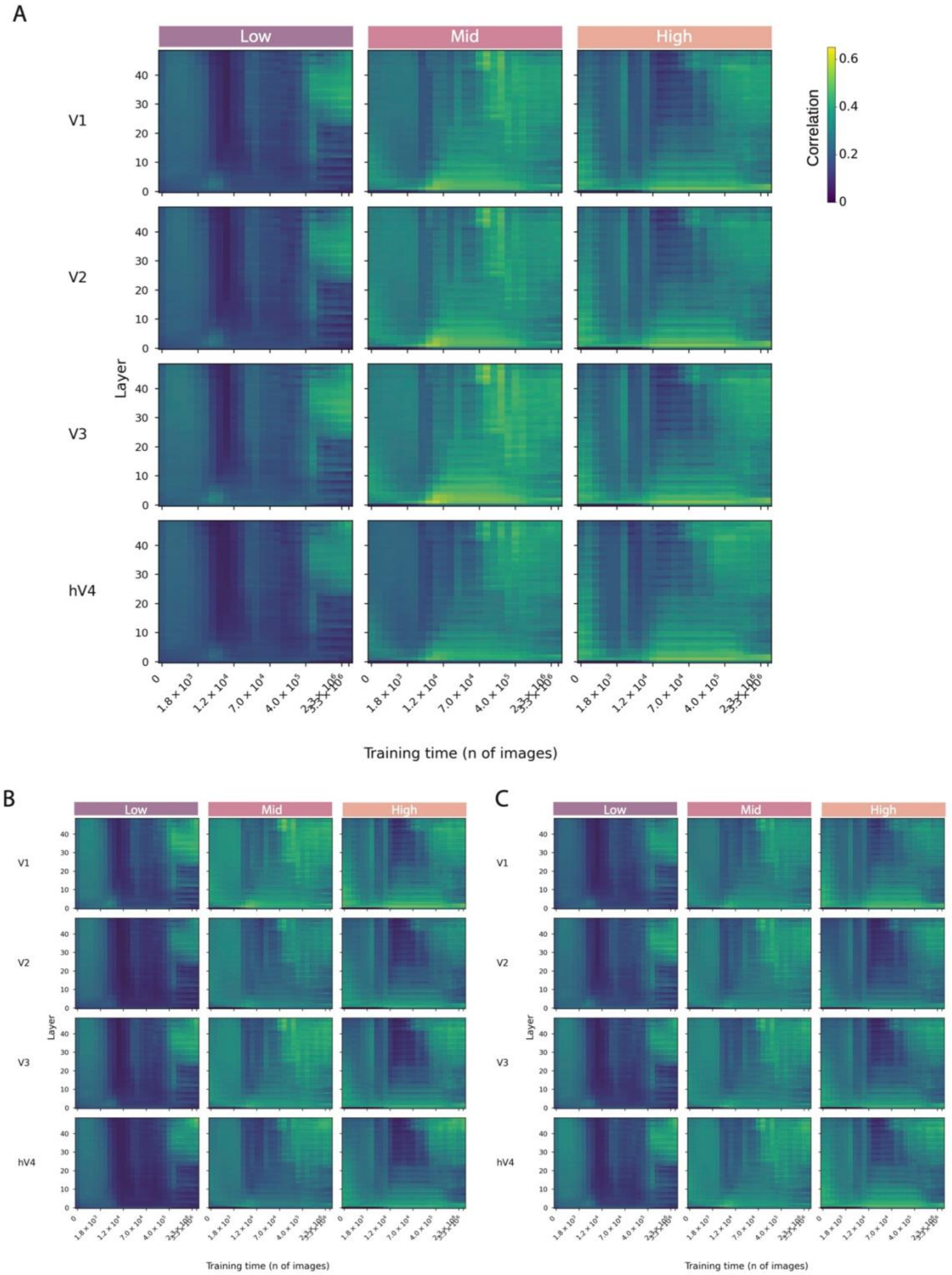


**Figure S17. Layer-wise trajectories of model–brain correspondence across the critical training window.**

Layer-wise model–brain correspondence across the critical training window for low-, mid-, and high-spatial-frequency models in adults (A), 9-month-olds (B), and 2-month-olds (C). Heat-map color indicates the bootstrap median correlation, normalized by the age- and ROI-specific noise ceiling, for each network layer and sampled training checkpoint after averaging across the 10 independently initialized network instances.

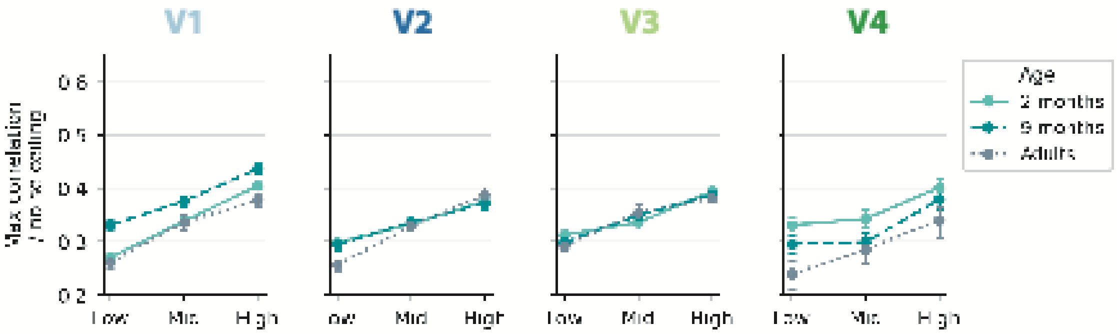


**Figure S18. Band-specific model–brain correspondence before object training.**
Peak-layer model–brain correspondence for representations derived from low-, mid-, and high-spatial-frequency bandpass-filtered FOUNDCOG stimuli in the untrained network across V1, V2, V3, and V4 in 2-month-olds, 9-month-olds, and adults. Model–brain correspondence was normalized by the age- and ROI-specific noise ceiling. Points show bootstrap medians and error bars show 95% bootstrap percentile intervals.

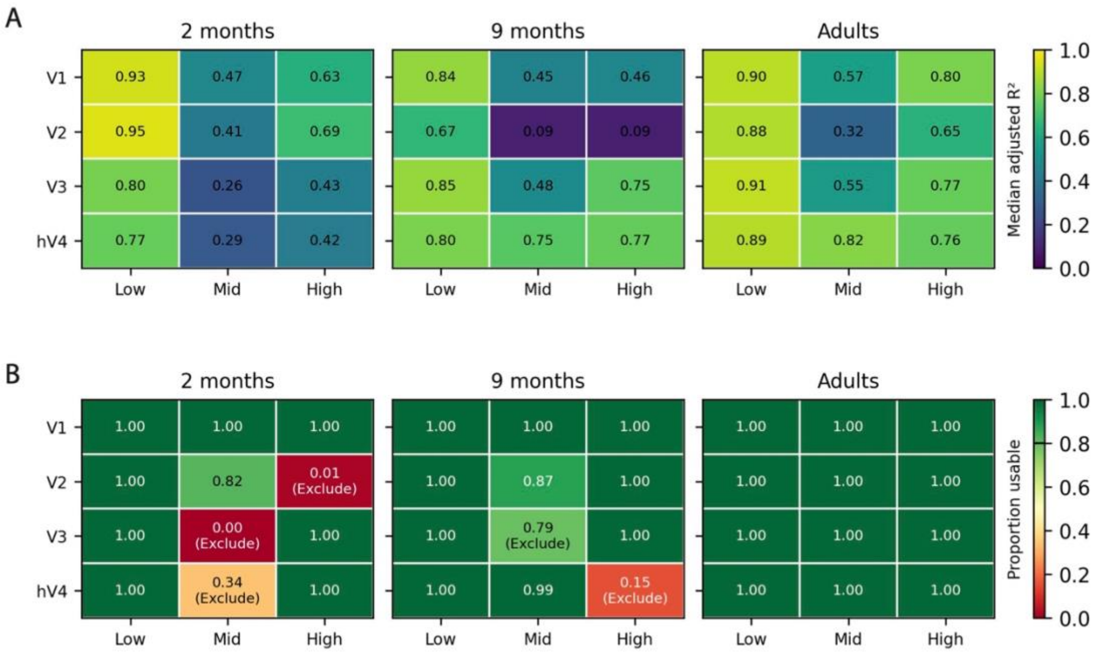


**Figure S19. Evaluation of logistic fit quality across ages and visual regions during the critical training window.**

(A) Median adjusted coefficient of determination ($R^2_{\text{adj}}$) across bootstrap logistic fits for each age × ROI × spatial-frequency-band combination. (B) Proportion of bootstrap fits meeting the specified usability criteria. Age × ROI × band combinations were considered sufficiently reliable for interpretation when at least 80% of bootstrap fits were usable. Combinations failing this criterion are marked 'Exclude'. In the primary analysis, 31 of 36 combinations met the reliability criterion, compared with 25 of 36 in the preliminary single-model analysis used to define the critical training window (Figure S16).

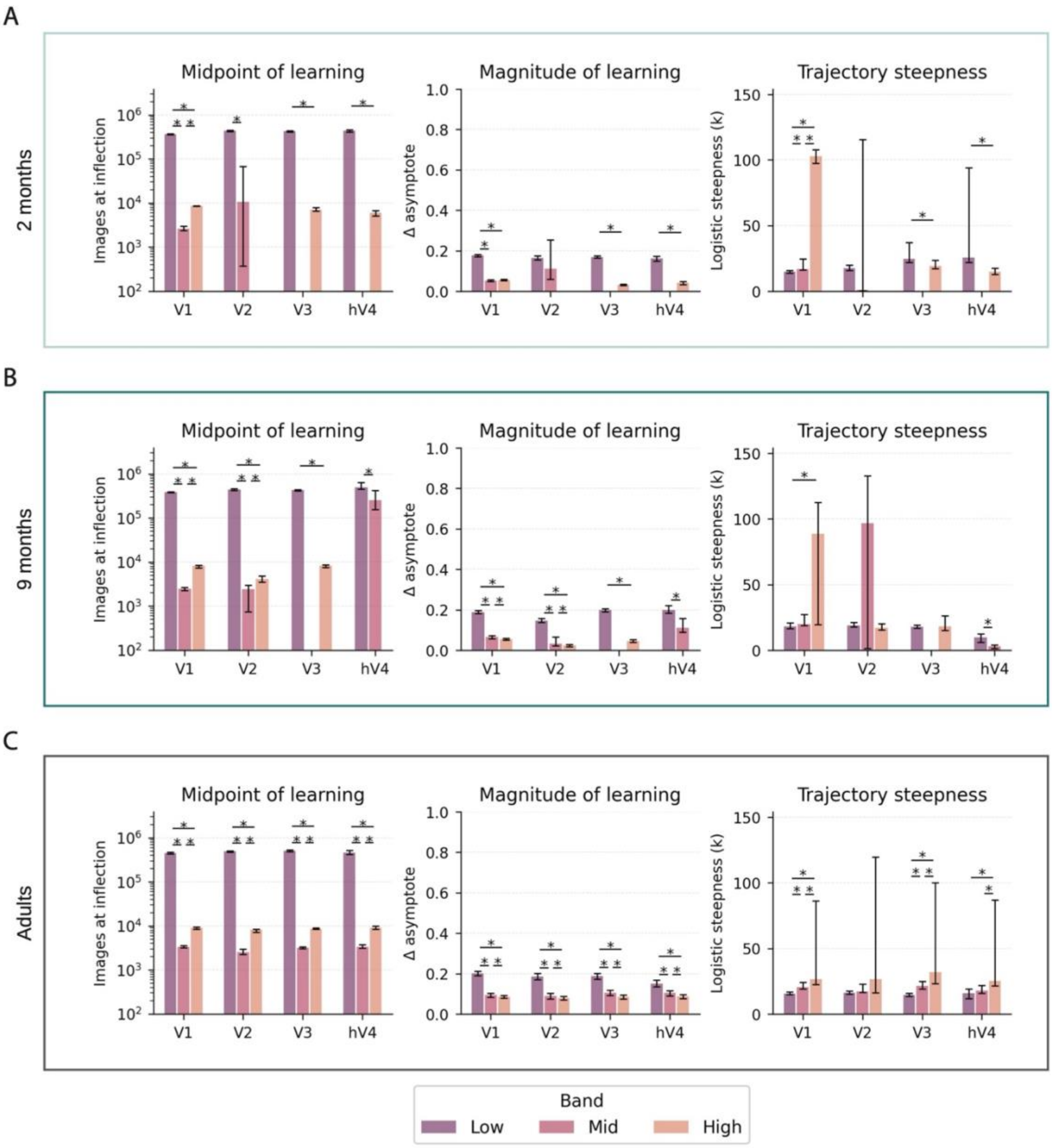


**Figure S20. Band-specific differences in logistic training dynamics across visual regions and ages.** Bootstrap estimates of the three logistic parameters used to characterize learning dynamics (midpoint of learning, magnitude of learning, and trajectory steepness) across low-, mid-, and high-spatial-frequency models and visual regions in 2-month-olds (A), 9-month-olds (B), and adults (C). Only age × ROI × band combinations meeting the fit-reliability criterion were included.

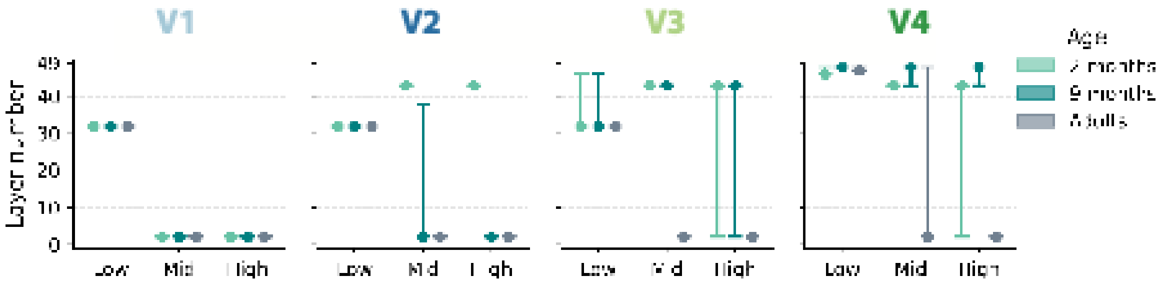


**Figure S21. Peak-corresponding DNN layers for spatial-frequency-specific representations at the end of the critical training window.**
Low-band correspondence peaked at a consistent depth across ages, in intermediate layers in V1–V3 and in late layers in V4. Mid- and high-band correspondence peaked in the earliest model layers in V1 at all ages and in adult V2–V4, but in later layers in infant V3 and V4 at both ages and in V2 at 2 months. For each spatial-frequency band, age group, and ROI, layer-wise model–brain correlations were averaged across the 10 independently initialized network instances, and the peak-corresponding layer was defined as the layer yielding the highest correspondence. Points show bootstrap medians and error bars show 95% bootstrap percentile intervals; wide intervals reflect bootstrap samples in which the peak alternated between early and late layers. Corresponding layer-wise model–brain correspondence across the critical training window is shown in Figure S17.

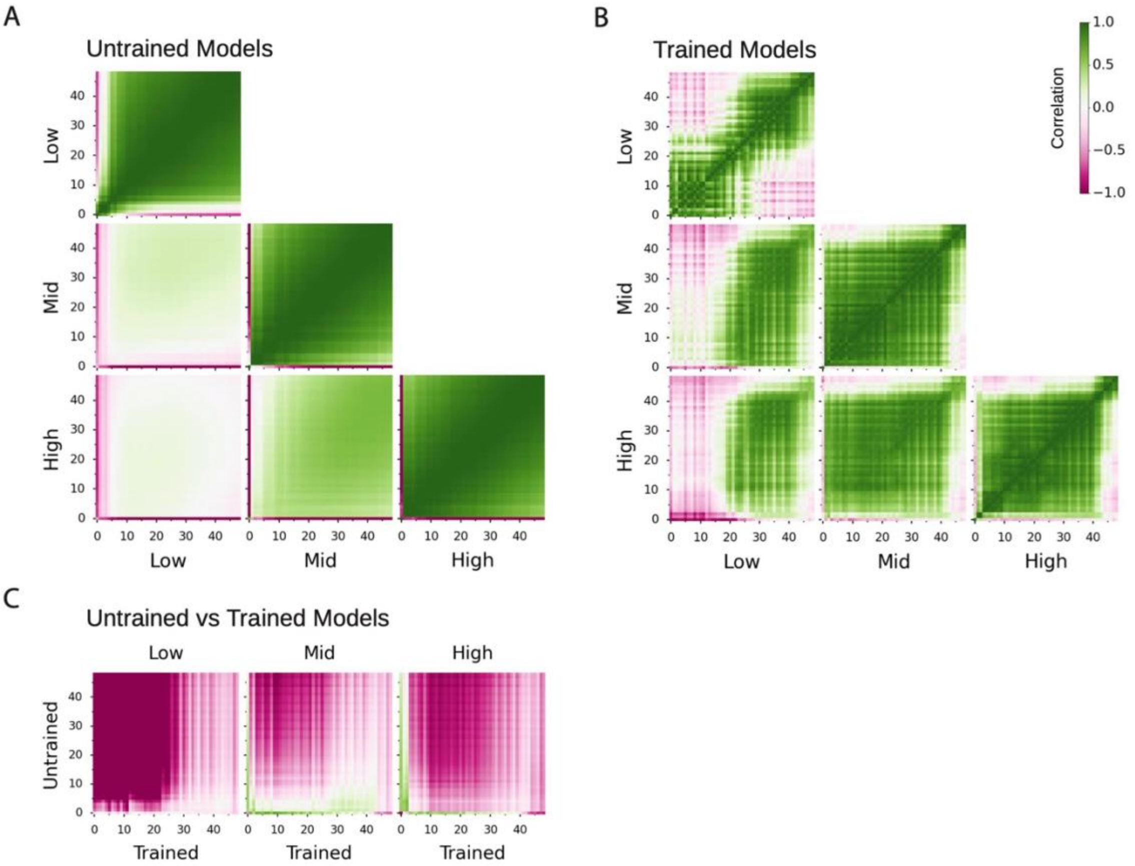


**Figure S22. Convergence of model representations derived from visual information at different spatial scales before and after the critical training window.**
Layer-wise representational similarity across low-, mid-, and high-spatial-frequency models, averaged across 10 independently initialized network instances per band. (A) Layer-wise similarity within and between bands in untrained models. (B) The same comparisons at the end of the critical training window. (C) Layer-wise similarity between untrained and trained representations within each band. Heat-map color indicates Spearman correlation.

### Supplemental Tables

**Table S1.** Split-half noise ceilings for regions of interest along the ventral visual stream.

| | 2 months | 9 months | Adult |
|---|---|---|---|
| **V1** | 0.9422 | 0.9383 | 0.9516 |
| **V2dv** | 0.9010 | 0.9077 | 0.9399 |
| **V3v** | 0.9463 | 0.9432 | 0.9458 |
| **hV4** | 0.8495 | 0.8569 | 0.8488 |
| **VO1** | 0.6954 | 0.7460 | 0.6618 |
| **FG4** | 0.3211 | 0.6016 | 0.7263 |

**Table S2.** Bootstrapped pairwise differences in peak-layer model–brain correspondence between spatial-frequency bands in the untrained model (averaged across 10 independently initialized network instances).

| **ROI** | **Age** | **Comparison** | **Median paired difference** | **Significant** |
|---|---|---|---|---|
| V1 | 2 months | Mid minus Low | 0.068 | Yes |
| V1 | 2 months | High minus Low | 0.135 | Yes |
| V1 | 2 months | High minus Mid | 0.067 | Yes |
| V1 | 9 months | Mid minus Low | 0.045 | Yes |
| V1 | 9 months | High minus Low | 0.105 | Yes |
| V1 | 9 months | High minus Mid | 0.06 | Yes |
| V1 | Adult | Mid minus Low | 0.075 | Yes |
| V1 | Adult | High minus Low | 0.117 | Yes |
| V1 | Adult | High minus Mid | 0.042 | Yes |
| V2dv | 2 months | Mid minus Low | 0.034 | Yes |
| V2dv | 2 months | High minus Low | 0.078 | Yes |
| V2dv | 2 months | High minus Mid | 0.044 | Yes |
| V2dv | 9 months | Mid minus Low | 0.041 | Yes |
| V2dv | 9 months | High minus Low | 0.078 | Yes |
| V2dv | 9 months | High minus Mid | 0.037 | Yes |
| V2dv | Adult | Mid minus Low | 0.073 | Yes |
| V2dv | Adult | High minus Low | 0.131 | Yes |
| V2dv | Adult | High minus Mid | 0.059 | Yes |
| V3v | 2 months | Mid minus Low | 0.024 | Yes |
| V3v | 2 months | High minus Low | 0.082 | Yes |
| V3v | 2 months | High minus Mid | 0.059 | Yes |
| V3v | 9 months | Mid minus Low | 0.05 | Yes |
| V3v | 9 months | High minus Low | 0.089 | Yes |
| V3v | 9 months | High minus Mid | 0.039 | Yes |
| V3v | Adult | Mid minus Low | 0.062 | Yes |
| V3v | Adult | High minus Low | 0.089 | Yes |
| V3v | Adult | High minus Mid | 0.026 | Yes |
| hV4 | 2 months | Mid minus Low | 0.012 | No |
| hV4 | 2 months | High minus Low | 0.071 | Yes |
| hV4 | 2 months | High minus Mid | 0.059 | Yes |
| hV4 | 9 months | Mid minus Low | 0.004 | No |
| hV4 | 9 months | High minus Low | 0.084 | Yes |
| hV4 | 9 months | High minus Mid | 0.08 | Yes |
| hV4 | Adult | Mid minus Low | 0.045 | Yes |
| hV4 | Adult | High minus Low | 0.099 | Yes |
| hV4 | Adult | High minus Mid | 0.055 | Yes |

**Table S3.** Bootstrapped changes in peak-layer model–brain correspondence from the untrained model to the end of the critical training window for each spatial-frequency band (averaged across 10 independently initialized network instances).

| Age | ROI | Band | Median difference | 95% bootstrap CI | Significant increase |
|---|---|---|---|---|---|
| 2 months | V1 | Low | 0.132 | [0.1246, 0.1392] | Yes |
| 2 months | V1 | Mid | 0.0776 | [0.0711, 0.0833] | Yes |
| 2 months | V1 | High | 0.0723 | [0.0672, 0.0768] | Yes |
| 2 months | V2 | Low | 0.1355 | [0.1238, 0.1477] | Yes |
| 2 months | V2 | Mid | 0.0654 | [0.0539, 0.0770] | Yes |
| 2 months | V2 | High | 0.045 | [0.0320, 0.0576] | Yes |
| 2 months | V3 | Low | 0.0991 | [0.0916, 0.1070] | Yes |
| 2 months | V3 | Mid | 0.0881 | [0.0810, 0.0957] | Yes |
| 2 months | V3 | High | 0.0485 | [0.0424, 0.0564] | Yes |
| 2 months | hV4 | Low | 0.0989 | [0.0829, 0.1150] | Yes |
| 2 months | hV4 | Mid | 0.0919 | [0.0764, 0.1065] | Yes |
| 2 months | hV4 | High | 0.0679 | [0.0525, 0.0845] | Yes |
| 9 months | V1 | Low | 0.1334 | [0.1218, 0.1437] | Yes |
| 9 months | V1 | Mid | 0.0853 | [0.0740, 0.0952] | Yes |
| 9 months | V1 | High | 0.0795 | [0.0717, 0.0871] | Yes |
| 9 months | V2 | Low | 0.0676 | [0.0536, 0.0813] | Yes |
| 9 months | V2 | Mid | 0.0157 | [0.0035, 0.0284] | Yes |
| 9 months | V2 | High | 0.0531 | [0.0445, 0.0610] | Yes |
| 9 months | V3 | Low | 0.1329 | [0.1209, 0.1445] | Yes |
| 9 months | V3 | Mid | 0.079 | [0.0689, 0.0888] | Yes |
| 9 months | V3 | High | 0.0696 | [0.0580, 0.0810] | Yes |
| 9 months | hV4 | Low | 0.1488 | [0.1290, 0.1682] | Yes |
| 9 months | hV4 | Mid | 0.1129 | [0.0924, 0.1344] | Yes |
| 9 months | hV4 | High | 0.1034 | [0.0806, 0.1267] | Yes |
| Adults | V1 | Low | 0.1865 | [0.1711, 0.2015] | Yes |
| Adults | V1 | Mid | 0.109 | [0.0979, 0.1192] | Yes |
| Adults | V1 | High | 0.1246 | [0.1125, 0.1348] | Yes |
| Adults | V2 | Low | 0.1739 | [0.1552, 0.1926] | Yes |
| Adults | V2 | Mid | 0.098 | [0.0803, 0.1152] | Yes |
| Adults | V2 | High | 0.1258 | [0.1120, 0.1383] | Yes |
| Adults | V3 | Low | 0.1712 | [0.1487, 0.1922] | Yes |
| Adults | V3 | Mid | 0.1175 | [0.1029, 0.1304] | Yes |
| Adults | V3 | High | 0.1351 | [0.1182, 0.1502] | Yes |
| Adults | hV4 | Low | 0.1461 | [0.1193, 0.1694] | Yes |
| Adults | hV4 | Mid | 0.1288 | [0.1154, 0.1410] | Yes |
| Adults | hV4 | High | 0.1448 | [0.1295, 0.1583] | Yes |

**Table S4.** Spatial-frequency-band differences in the dynamics of model–brain correspondence throughout training across ages and visual regions.

| Metric | Test | Comparison | n | df | Statistic | p-value | p-value Holm corrected | Median difference | Significant |
|---|---|---|---|---|---|---|---|---|---|
| Midpoint of learning | Skillings-Mack | Omnibus | 12 | 2 | 18.5143 | <0.001 | – | – | Yes |
| Midpoint of learning | Wilcoxon signed rank | Mid minus Low | 9 | – | 0 | 0.0039 | 0.0078 | -444454 | Yes |
| Midpoint of learning | Wilcoxon signed rank | High minus Low | 10 | – | 0 | 0.002 | 0.0059 | -438219 | Yes |
| Midpoint of learning | Wilcoxon signed rank | High minus Mid | 7 | – | 0 | 0.0156 | 0.0156 | 5798 | Yes |
| Magnitude of learning | Skillings-Mack | Omnibus | 12 | 2 | 17.068 | <0.001 | – | – | Yes |
| Magnitude of learning | Wilcoxon signed rank | Mid minus Low | 9 | – | 0 | 0.0039 | 0.0078 | -0.0975 | Yes |
| Magnitude of learning | Wilcoxon signed rank | High minus Low | 10 | – | 0 | 0.002 | 0.0059 | -0.1204 | Yes |
| Magnitude of learning | Wilcoxon signed rank | High minus Mid | 7 | – | 1 | 0.0312 | 0.0312 | -0.0107 | Yes |
| Trajectory steepness | Skillings-Mack | Omnibus | 12 | 2 | 4.0868 | 0.1296 | – | – | No |
| Trajectory steepness | Wilcoxon signed rank | Mid minus Low | 9 | – | 14 | 0.3594 | 0.4375 | 2.1484 | No |
| Trajectory steepness | Wilcoxon signed rank | High minus Low | 10 | – | 11 | 0.1055 | 0.3164 | 9.9389 | No |
| Trajectory steepness | Wilcoxon signed rank | High minus Mid | 7 | – | 6 | 0.2188 | 0.4375 | 9.4718 | No |

**Table S5.** Age- and region-specific pairwise differences in the dynamics of model–brain correspondence across spatial-frequency bands throughout training.

| Age | ROI | Metric | Comparison | Median difference | 95% CI | Significant |
|---|---|---|---|---|---|---|
| 2 months | V1 | Midpoint of learning | Mid minus Low | -365529 | [-375633, -355356] | Yes |
| 2 months | V1 | Midpoint of learning | High minus Low | -359541 | [-369581, -349482] | Yes |
| 2 months | V1 | Midpoint of learning | High minus Mid | 5997 | [5595, 6190] | Yes |
| 2 months | V2 | Midpoint of learning | Mid minus Low | -426278 | [-449981, -359258] | Yes |
| 2 months | V2 | Midpoint of learning | High minus Low | – | – | – |
| 2 months | V2 | Midpoint of learning | High minus Mid | – | – | – |
| 2 months | V3 | Midpoint of learning | Mid minus Low | – | – | – |
| 2 months | V3 | Midpoint of learning | High minus Low | -425588 | [-438195, -409334] | Yes |
| 2 months | V3 | Midpoint of learning | High minus Mid | – | – | – |
| 2 months | hV4 | Midpoint of learning | Mid minus Low | – | – | – |
| 2 months | hV4 | Midpoint of learning | High minus Low | -433499 | [-458459, -403654] | Yes |
| 2 months | hV4 | Midpoint of learning | High minus Mid | – | – | – |
| 2 months | V1 | Magnitude of learning | Mid minus Low | -0.123 | [-0.130, -0.118] | Yes |
| 2 months | V1 | Magnitude of learning | High minus Low | -0.121 | [-0.126, -0.116] | Yes |
| 2 months | V1 | Magnitude of learning | High minus Mid | 0.003 | [-0.001, 0.006] | No |
| 2 months | V2 | Magnitude of learning | Mid minus Low | -0.054 | [-0.101, 0.086] | No |
| 2 months | V2 | Magnitude of learning | High minus Low | – | – | – |
| 2 months | V2 | Magnitude of learning | High minus Mid | – | – | – |
| 2 months | V3 | Magnitude of learning | Mid minus Low | – | – | – |
| 2 months | V3 | Magnitude of learning | High minus Low | -0.137 | [-0.143, -0.132] | Yes |
| 2 months | V3 | Magnitude of learning | High minus Mid | – | – | – |
| 2 months | hV4 | Magnitude of learning | Mid minus Low | – | – | – |
| 2 months | hV4 | Magnitude of learning | High minus Low | -0.12 | [-0.131, -0.107] | Yes |

| | | | | | | |
|---|---|---|---|---|---|---|
| 2 months | hV4 | Magnitude of learning | High minus Mid | – | – | – |
| 2 months | V1 | Trajectory steepness | Mid minus Low | 2.191 | [0.854, 10.823] | Yes |
| 2 months | V1 | Trajectory steepness | High minus Low | 88.062 | [82.490, 93.124] | Yes |
| 2 months | V1 | Trajectory steepness | High minus Mid | 85.669 | [73.009, 90.790] | Yes |
| 2 months | V2 | Trajectory steepness | Mid minus Low | -16.515 | [-19.056, 97.739] | No |
| 2 months | V2 | Trajectory steepness | High minus Low | – | – | – |
| 2 months | V2 | Trajectory steepness | High minus Mid | – | – | – |
| 2 months | V3 | Trajectory steepness | Mid minus Low | – | – | – |
| 2 months | V3 | Trajectory steepness | High minus Low | -5.588 | [-17.658, -0.206] | Yes |
| 2 months | V3 | Trajectory steepness | High minus Mid | – | – | – |
| 2 months | hV4 | Trajectory steepness | Mid minus Low | – | – | – |
| 2 months | hV4 | Trajectory steepness | High minus Low | -11.091 | [-79.683, -6.121] | Yes |
| 2 months | hV4 | Trajectory steepness | High minus Mid | – | – | – |
| 9 months | V1 | Midpoint of learning | Mid minus Low | -386602 | [-395573, -375270] | Yes |
| 9 months | V1 | Midpoint of learning | High minus Low | -380976 | [-389951, -369593] | Yes |
| 9 months | V1 | Midpoint of learning | High minus Mid | 5776 | [4976, 6020] | Yes |
| 9 months | V2 | Midpoint of learning | Mid minus Low | -444616 | [-464004, -426157] | Yes |
| 9 months | V2 | Midpoint of learning | High minus Low | -442757 | [-462313, -423800] | Yes |
| 9 months | V2 | Midpoint of learning | High minus Mid | 1737 | [1078, 3722] | Yes |
| 9 months | V3 | Midpoint of learning | Mid minus Low | – | – | – |
| 9 months | V3 | Midpoint of learning | High minus Low | -420826 | [-433047, -409480] | Yes |
| 9 months | V3 | Midpoint of learning | High minus Mid | – | – | – |
| 9 months | hV4 | Midpoint of learning | Mid minus Low | -258277 | [-388091, -84459] | Yes |
| 9 months | hV4 | Midpoint of learning | High minus Low | – | – | – |
| 9 months | hV4 | Midpoint of learning | High minus Mid | – | – | – |

| | | | | | | |
|---|---|---|---|---|---|---|
| 9 months | V1 | Magnitude of learning | Mid minus Low | -0.124 | [-0.133, -0.116] | Yes |
| 9 months | V1 | Magnitude of learning | High minus Low | -0.135 | [-0.142, -0.127] | Yes |
| 9 months | V1 | Magnitude of learning | High minus Mid | -0.011 | [-0.015, -0.006] | Yes |
| 9 months | V2 | Magnitude of learning | Mid minus Low | -0.114 | [-0.125, -0.080] | Yes |
| 9 months | V2 | Magnitude of learning | High minus Low | -0.124 | [-0.132, -0.116] | Yes |
| 9 months | V2 | Magnitude of learning | High minus Mid | -0.009 | [-0.043, -0.003] | Yes |
| 9 months | V3 | Magnitude of learning | Mid minus Low | – | – | – |
| 9 months | V3 | Magnitude of learning | High minus Low | -0.152 | [-0.159, -0.145] | Yes |
| 9 months | V3 | Magnitude of learning | High minus Mid | – | – | – |
| 9 months | hV4 | Magnitude of learning | Mid minus Low | -0.088 | [-0.108, -0.044] | Yes |
| 9 months | hV4 | Magnitude of learning | High minus Low | – | – | – |
| 9 months | hV4 | Magnitude of learning | High minus Mid | – | – | – |
| 9 months | V1 | Trajectory steepness | Mid minus Low | 1.693 | [-1.026, 7.512] | No |
| 9 months | V1 | Trajectory steepness | High minus Low | 70.149 | [0.421, 94.356] | Yes |
| 9 months | V1 | Trajectory steepness | High minus Mid | 68.559 | [-6.423, 93.580] | No |
| 9 months | V2 | Trajectory steepness | Mid minus Low | 77.453 | [-17.573, 114.182] | No |
| 9 months | V2 | Trajectory steepness | High minus Low | -1.946 | [-4.463, 1.440] | No |
| 9 months | V2 | Trajectory steepness | High minus Mid | -80.135 | [-114.384, 14.905] | No |
| 9 months | V3 | Trajectory steepness | Mid minus Low | – | – | – |
| 9 months | V3 | Trajectory steepness | High minus Low | 0.188 | [-2.998, 8.535] | No |
| 9 months | V3 | Trajectory steepness | High minus Mid | – | – | – |
| 9 months | hV4 | Trajectory steepness | Mid minus Low | -6.598 | [-10.030, -2.714] | Yes |
| 9 months | hV4 | Trajectory steepness | High minus Low | – | – | – |
| 9 months | hV4 | Trajectory steepness | High minus Mid | – | – | – |
| Adults | V1 | Midpoint of learning | Mid minus Low | -450043 | [-468293, -433623] | Yes |

| | | | | | | |
|---|---|---|---|---|---|---|
| Adults | V1 | Midpoint of learning | High minus Low | -444405 | [-462468, -427869] | Yes |
| Adults | V1 | Midpoint of learning | High minus Mid | 5665 | [5392, 5889] | Yes |
| Adults | V2 | Midpoint of learning | Mid minus Low | -489634 | [-510246, -470049] | Yes |
| Adults | V2 | Midpoint of learning | High minus Low | -483966 | [-504463, -464361] | Yes |
| Adults | V2 | Midpoint of learning | High minus Mid | 5672 | [4963, 5969] | Yes |
| Adults | V3 | Midpoint of learning | Mid minus Low | -503922 | [-524314, -484023] | Yes |
| Adults | V3 | Midpoint of learning | High minus Low | -498291 | [-518683, -478547] | Yes |
| Adults | V3 | Midpoint of learning | High minus Mid | 5536 | [5355, 5803] | Yes |
| Adults | hV4 | Midpoint of learning | Mid minus Low | -456063 | [-518746, -418229] | Yes |
| Adults | hV4 | Midpoint of learning | High minus Low | -450225 | [-513088, -412020] | Yes |
| Adults | hV4 | Midpoint of learning | High minus Mid | 5806 | [5421, 6339] | Yes |
| Adults | V1 | Magnitude of learning | Mid minus Low | -0.107 | [-0.117, -0.096] | Yes |
| Adults | V1 | Magnitude of learning | High minus Low | -0.115 | [-0.125, -0.106] | Yes |
| Adults | V1 | Magnitude of learning | High minus Mid | -0.009 | [-0.014, -0.004] | Yes |
| Adults | V2 | Magnitude of learning | Mid minus Low | -0.098 | [-0.113, -0.082] | Yes |
| Adults | V2 | Magnitude of learning | High minus Low | -0.108 | [-0.120, -0.096] | Yes |
| Adults | V2 | Magnitude of learning | High minus Mid | -0.011 | [-0.017, -0.003] | Yes |
| Adults | V3 | Magnitude of learning | Mid minus Low | -0.083 | [-0.096, -0.069] | Yes |
| Adults | V3 | Magnitude of learning | High minus Low | -0.104 | [-0.115, -0.091] | Yes |
| Adults | V3 | Magnitude of learning | High minus Mid | -0.021 | [-0.026, -0.015] | Yes |
| Adults | hV4 | Magnitude of learning | Mid minus Low | -0.046 | [-0.068, -0.027] | Yes |
| Adults | hV4 | Magnitude of learning | High minus Low | -0.064 | [-0.084, -0.046] | Yes |
| Adults | hV4 | Magnitude of learning | High minus Mid | -0.018 | [-0.029, -0.006] | Yes |
| Adults | V1 | Trajectory steepness | Mid minus Low | 5.098 | [3.662, 8.432] | Yes |
| Adults | V1 | Trajectory steepness | High minus Low | 11.027 | [7.109, 69.362] | Yes |

| | | | | | | |
|---|---|---|---|---|---|---|
| Adults | V1 | Trajectory steepness | High minus Mid | 5.899 | [2.854, 60.713] | Yes |
| Adults | V2 | Trajectory steepness | Mid minus Low | 1.022 | [-0.143, 7.298] | No |
| Adults | V2 | Trajectory steepness | High minus Low | 10.417 | [-0.891, 103.346] | No |
| Adults | V2 | Trajectory steepness | High minus Mid | 9.729 | [-4.547, 102.772] | No |
| Adults | V3 | Trajectory steepness | Mid minus Low | 6.885 | [4.680, 10.092] | Yes |
| Adults | V3 | Trajectory steepness | High minus Low | 17.385 | [9.157, 85.440] | Yes |
| Adults | V3 | Trajectory steepness | High minus Mid | 10.379 | [3.847, 77.251] | Yes |
| Adults | hV4 | Trajectory steepness | Mid minus Low | 2.514 | [-1.104, 7.345] | No |
| Adults | hV4 | Trajectory steepness | High minus Low | 9.631 | [4.407, 71.171] | Yes |
| Adults | hV4 | Trajectory steepness | High minus Mid | 7.254 | [3.985, 65.875] | Yes |

**Table S6.** Bootstrapped pairwise differences in peak-layer model–brain correspondence between spatial-frequency bands at the end of the critical training window (averaged across 10 independently initialized network instances).

| ROI | Age | Comparison | Median paired difference | Significant |
|---|---|---|---|---|
| V1 | 2 months | Mid minus Low | 0.01 | Yes |
| V1 | 2 months | High minus Low | 0.071 | Yes |
| V1 | 2 months | High minus Mid | 0.061 | Yes |
| V1 | 9 months | Mid minus Low | -0.006 | No |
| V1 | 9 months | High minus Low | 0.048 | Yes |
| V1 | 9 months | High minus Mid | 0.054 | Yes |
| V1 | Adult | Mid minus Low | -0.006 | No |
| V1 | Adult | High minus Low | 0.051 | Yes |
| V1 | Adult | High minus Mid | 0.057 | Yes |
| V2dv | 2 months | Mid minus Low | -0.044 | Yes |
| V2dv | 2 months | High minus Low | -0.023 | Yes |
| V2dv | 2 months | High minus Mid | 0.021 | Yes |
| V2dv | 9 months | Mid minus Low | -0.016 | Yes |
| V2dv | 9 months | High minus Low | 0.061 | Yes |
| V2dv | 9 months | High minus Mid | 0.078 | Yes |
| V2dv | Adult | Mid minus Low | -0.008 | No |
| V2dv | Adult | High minus Low | 0.08 | Yes |
| V2dv | Adult | High minus Mid | 0.088 | Yes |
| V3v | 2 months | Mid minus Low | 0.012 | Yes |
| V3v | 2 months | High minus Low | 0.029 | Yes |
| V3v | 2 months | High minus Mid | 0.017 | Yes |
| V3v | 9 months | Mid minus Low | -0.007 | Yes |
| V3v | 9 months | High minus Low | 0.022 | Yes |
| V3v | 9 months | High minus Mid | 0.029 | Yes |
| V3v | Adult | Mid minus Low | 0.006 | No |
| V3v | Adult | High minus Low | 0.051 | Yes |
| V3v | Adult | High minus Mid | 0.045 | Yes |
| hV4 | 2 months | Mid minus Low | 0.004 | No |
| hV4 | 2 months | High minus Low | 0.034 | Yes |
| hV4 | 2 months | High minus Mid | 0.031 | Yes |
| hV4 | 9 months | Mid minus Low | -0.038 | Yes |
| hV4 | 9 months | High minus Low | 0.031 | Yes |
| hV4 | 9 months | High minus Mid | 0.068 | Yes |
| hV4 | Adult | Mid minus Low | 0.025 | No |
| hV4 | Adult | High minus Low | 0.098 | Yes |
| hV4 | Adult | High minus Mid | 0.073 | Yes |

**Table S7.** Adult–infant differences in relative band-specific peak-layer model–brain correspondence at the end of the critical training window across visual regions.

| Comparison | ROI | Band | Median difference (Fisher z) | 95% bootstrap CI | Significant |
|---|---|---|---|---|---|
| Adult minus 2 months | V1 | Low | 0.0128 | [0.0015, 0.0238] | Yes |
| Adult minus 9 months | V1 | Low | -0.0015 | [-0.0128, 0.0102] | No |
| Adult minus 2 months | V1 | Mid | -0.0056 | [-0.0129, 0.0012] | No |
| Adult minus 9 months | V1 | Mid | -0.0012 | [-0.0093, 0.0068] | No |
| Adult minus 2 months | V1 | High | -0.0072 | [-0.0178, 0.0041] | No |
| Adult minus 9 months | V1 | High | 0.0026 | [-0.0092, 0.0144] | No |
| Adult minus 2 months | V2 | Low | -0.0518 | [-0.0625, -0.0403] | Yes |
| Adult minus 9 months | V2 | Low | -0.0124 | [-0.0230, -0.0002] | Yes |
| Adult minus 2 months | V2 | Mid | -0.015 | [-0.0238, -0.0062] | Yes |
| Adult minus 9 months | V2 | Mid | -0.0056 | [-0.0148, 0.0034] | No |
| Adult minus 2 months | V2 | High | 0.067 | [0.0557, 0.0772] | Yes |
| Adult minus 9 months | V2 | High | 0.0177 | [0.0056, 0.0292] | Yes |
| Adult minus 2 months | V3 | Low | -0.0072 | [-0.0208, 0.0068] | No |
| Adult minus 9 months | V3 | Low | -0.017 | [-0.0302, -0.0023] | Yes |
| Adult minus 2 months | V3 | Mid | -0.0141 | [-0.0220, -0.0065] | Yes |
| Adult minus 9 months | V3 | Mid | -0.0025 | [-0.0104, 0.0058] | No |
| Adult minus 2 months | V3 | High | 0.0214 | [0.0095, 0.0331] | Yes |
| Adult minus 9 months | V3 | High | 0.0194 | [0.0075, 0.0303] | Yes |
| Adult minus 2 months | hV4 | Low | -0.0284 | [-0.0500, -0.0074] | Yes |
| Adult minus 9 months | hV4 | Low | -0.0424 | [-0.0651, -0.0216] | Yes |
| Adult minus 2 months | hV4 | Mid | -0.0074 | [-0.0200, 0.0054] | No |
| Adult minus 9 months | hV4 | Mid | 0.0197 | [0.0062, 0.0328] | Yes |
| Adult minus 2 months | hV4 | High | 0.0359 | [0.0183, 0.0524] | Yes |
| Adult minus 9 months | hV4 | High | 0.0238 | [0.0055, 0.0417] | Yes |
